# Progranulin promotes immune evasion of pancreatic adenocarcinoma through regulation of MHCI expression

**DOI:** 10.1101/2021.02.24.432700

**Authors:** Phyllis F Cheung, JiaJin Yang, Kirsten Krengel, Kristina Althoff, Chi Wai Yip, Elaine HL Siu, Linda WC Ng, Karl S Lang, Lamin Cham, Daniel R Engel, Camille Soun, Igor Cima, Björn Scheffler, Jana K Striefler, Marianne Sinn, Marcus Bahra, Uwe Pelzer, Helmut Oettle, Peter Markus, Esther MM Smeets, Erik HJG Aarntzen, Konstantinos Savvatakis, Sven-Thorsten Liffers, Christian Neander, Anna Bazarna, Xin Zhang, Doron Merkler, Annette Paschen, Howard C. Crawford, Anthony WH Chan, Siu Tim Cheung, Jens T Siveke

## Abstract

Immune evasion is indispensable for cancer initiation and progression, although its underlying mechanisms in pancreatic ductal adenocarcinoma (PDAC) remain elusive. Here, we unveiled a cancer cell-autonomous function of PGRN in driving immune evasion in primary PDAC. Tumor- but not macrophage-derived PGRN was associated with poor overall survival in PDAC. Multiplex immunohistochemistry revealed low MHC class I (MHCI) expression and lack of CD8+ T cells infiltration in PGRN-high tumors. Inhibition of PGRN abrogated autophagy-dependent MHCI degradation and restored MHCI expression on PDAC cells. Antibody-based blockade of PGRN in a genetic PDAC mouse model remarkably decelerated tumor initiation and progression. Notably, tumors expressing LCMV-gp33 as model antigen were sensitized towards cytotoxic gp33-TCR transgenic T cells upon anti-PGRN antibody treatment. Overall, our study uncovered an unprecedented role of tumor-derived PGRN in regulating immunogenicity of primary PDAC.

**STATEMENT OF SIGNIFICANCE:** Immune evasion is a key property of PDAC, rendering it refractory to immunotherapy. Here we demonstrate that tumor-derived PGRN promotes autophagy-dependent MHCI degradation, while anti-PGRN increases intratumoral CD8 infiltration and blocks tumor progression. With recent advances in T cell-mediated approaches, PGRN represents a pivotal target to enhance tumor antigen-specific cytotoxicity.

## INTRODUCTION

Although immunotherapy is an established treatment strategy in many cancers, pancreatic ductal adenocarcinoma (PDAC) is so far refractory to immunomodulatory approaches and still remains largely untreatable. The ability of tumor cells to evade immune elimination is fundamental to tumor initiation, progression and therapy resistance (1, 2). Down-regulation of major histocompatibility complex class I (MHCI) is a common mechanism evolved by neoplastic cells to evade immune recognition and cytotoxicity (3, 4). Efficacy of immunotherapies was reported to depend on the expression levels of MHCI on tumor cells(5–7). Understanding the mechanism to restore tumor MHCI expression is therefore critical to induce antitumor immunity in this deadly disease.

Recently, Nielsen *et al* showed that macrophage-derived progranulin (protein: PGRN; gene: *GRN*) promoted liver metastasis of PDAC (8) and contributed to immunotherapy resistance in metastatic PDAC (9). However, the study did not address the role of PGRN in tumor cells. Tumorigenic role of PGRN is well-documented in various cancers, where it promotes cell proliferation, migration and chemoresistance (10–15). In HCC, PGRN enhances the shedding of tumor cell MHC class I chain-related protein A (MICA), an innate NK and T cell stimulatory molecule (16, 17), suggesting that PGRN might represent a tumor-intrinsic factor rendering tumor cells invisible to immune elimination. In PDAC, however, the role of tumor-derived PGRN in immune evasion is largely unaddressed.

Here, we applied multiplex immunohistochemistry and spatial analysis, as well as functional studies in human PDAC and next-generation genetic mouse models of spontaneous PDAC to elucidate the role of PGRN in immune evasion and tumor development of PDAC. Importantly, we used a spontaneous mouse model with inducible expression of lymphocytic choriomeningitis virus (LCMV)-glycoprotein (gp)33 as defined antigen to mechanistically address the effect of PGRN blockade in restoring tumor immunogenicity and tumor antigen-specific cytotoxicity.

## MATERIALS AND METHODS

### Clinical specimens

Expression of PGRN was analyzed in three independent cohorts of patients from the University Hospitals Essen (Essen cohort) and Radboud University Medical Center (Nijmegen cohort) and the phase III adjuvant CONKO-001 randomized trial (18).

For Essen cohort, a retrospective study was carried out according to the recommendations of the local ethics committee of the Medical Faculty of the University of Duisburg-Essen. Clinical data were obtained from archives and electronic health records. In this exploratory retrospective study, a cohort of 53 patients that had undergone pancreatic resection with a final histopathologic diagnosis of human PDAC between March 2006 and February 2016 was used (Approval no: 17-7340-BO).

Additionally, 31 patient samples from an independent cohort from Radboud University Medical Center, Nijmegen, were used to confirm the findings of Essen cohort. The Nijmegen cohort consisted of 31 patients with histologically proven pancreatic ductal adenocarcinoma (PDAC) between November 2004 and January 2015. Importantly, these patients underwent pancreatic resection and the whole tumor was enclosed in large format cassettes allowing to assess intratumoral heterogeneity. Given the retrospective nature of this study and the anonymized handling of data, informed consent was waived by the institutional review board (protocol CMO2018-4420).

For CONKO-001, the clinical details of this study have been described previously (18). In brief, 183 FFPE tissue samples of CONKO-001 patients were collected retrospectively. Tissues from 165 patients was suitable for tissue microarray (TMA) construction. To model the existence of intratumoral heterogeneity, three different tumor areas were selected for the construction of TMAs using a manual tissue microarrayer (Beecher Instruments, Wisconsin, USA). Here, we analyzed only the observation arm (n=71), in order to focus on the role of PGRN in PDAC without treatment intervention.

### Gene expression data analysis

Transcriptional profiling was performed by using dataset from Maurer *et al* (GSE93326), generated from 65 pairs of tumor epithelium and stroma laser capture microdissected samples and 15 bulk tumors (19). The raw count data of GSE93326 were normalized for library size into counts per million (CPM) by the RLE method implemented in EdgeR (3.28.0) to generate an expression table. We split the dataset into stroma and epithelium samples to analyze them separately. Both of them were further divided into *GRN*-high (n=32) and *GRN*-low (n=32) groups by the median CPM value of *GRN*. Differential expression profile was generated by EdgeR comparing the *GRN*-high and *GRN*-low groups. Gene set enrichment analysis (GSEA) was performed by fgsea (1.12.0) using the default parameters. Enrichment of the Hallmark and the KEGG pathways were tested against the differential expression profiles ranked by –log (p-value) × sign (log2FC). Pathways with the Benjamini-Hochberg method adjusted p-value (padj) smaller than 0.05 were considered significant.

### Cell type estimation using transcriptomes

Cell type-specific signals were determined similarly to Cima *et al* (20). First, we generated a reference map of 80 genes specific for 43 different cell types of interest using the primary cell atlas (GSE4991015) and the Q statistic introduced by Schug *et al* (21). Next, for each RNA-Seq query sample of the stroma RNA samples derived from the Maurer dataset (GSE93326), we selected genes with expression of >100 counts. From this data, we counted the occurrence of the specific genes for each cell type present in our reference map. To determine if the number of enriched genes was different from enrichment by chance, we compared the counts obtained using our reference map with 1000 randomly selected gene signatures. Finally, for each cell type in each experimental sample, a Fisher’s exact test was applied to determine whether the number of enriched specific genes was equal to the number of randomly enriched genes. The resulting odds ratios were used in inter-sample comparisons to generate hypotheses on the differential content of cell types present in bulk tissues **(Table S1).** To assess the association of each cell type with high and low *GRN* stromal samples, we computed 100x fold random forests for each cell type, similarly to Tan *et al* (22). The top 20 predictors for *GRN*-high and -low samples were selected and are presented in supplementary figure.

### Immunohistochemistry

Formalin-fixed paraffin-embedded (FFPE) sections were used for all IHC experiments. Antigen retrieval was performed by heat-induced epitope retrieval using citrate buffer (pH6), Tris/EDTA (pH9), or proteinase K treatment. After blocking with serum free protein blocking solution (Dako), slides were incubated for primary antibodies for 1h at RT, secondary antibody for 30 minutes at RT, and then subjected to Fast Red or DAB chromogen development. The details of antibodies are shown in **Table S2**. Slides were then counterstained with hematoxylin, dehydrated, and mounted. Stromal content and acinar cells were determined by Movat’s pentachrome staining following the manufacturer’s protocol (modified according to Verhoeff, Morphisto GmbH, Germany).

Slides were scanned and digitalized by Zeiss Axio Scanner Z.1 (Carl Zeiss AG, Germany) with 10x objective magnification. The percentage of positive cells for IHC staining was quantified by Definiens (Definiens AG, Germany). For quantification, total number of cells of the whole tissue section was determined based on nuclei staining (hematoxylin) detected by the software **(Figure S1)**.

### Multiplexed immunofluorescent (IF) histological staining

Multiplexed IF was performed using the Opal multiplex system (Perkin Elmer, MA) according to manufacturer’s instruction. In brief, FFPE sections were deparaffinized and then fixed with 4% paraformaldehyde prior to antigen retrieval by heat-induced epitope retrieval using citrate buffer (pH6) or Tris/EDTA (pH9). Each section was put through several sequential rounds of staining; each includes endogenous peroxidase blocking and non-specific protein blocking, followed by primary antibody and corresponding secondary horseradish peroxidase-conjugated polymer (Zytomed Systems, Germany or Perkin Elmer). Each horseradish peroxidase-conjugated polymer mediated the covalent binding of different fluorophore using tyramide signal amplification. Such covalent reaction was followed by additional antigen retrieval in heated citric buffer (pH6) or Tris/EDTA (pH9) for 10 min to remove antibodies before the next round of staining. After all sequential staining reactions, sections were counterstained with DAPI (Vector lab). The sequential multiplexed staining protocol is shown in **Table S3**. Slides were scanned and digitalized by Zeiss Axio Scanner Z.1 (Carl Zeiss AG, Germany) with 10x objective magnification.

### Spatial imaging analysis

An intensity threshold was used to generate masks for each fluorescent channel and the binary information for cellular and nuclear signals was coregistered. Automated analysis of cell distance was performed by a Java and R based algorithm. Using ImageJ, overlapping mask regions were employed to identify cells, which were marked with a point at the center of the DAPI^+^ cell nucleus.

The R Package Spatstat^1^ was used to convert cell coordinates to Euclidian distances between cells of interest. The algorithm determines the average distance and number of unique neighbors between any two cells. The threshold to pair up two cells was set to 50mm. The algorithm gives the percentage of MHCI+ cells in PGRN+/PanCK+ and PGRN-/PanCK+ populations, as well as the number of pairs between CD8+ and PGRN+/MHCI- or PGRN-/MHCI+ cells in each image(23).

### Mouse strains and tumor models

Animal experiments were approved under license number 84-02.04.2017.A315 by the Landesamt für Natur, Umwelt und Verbraucherschutz Nordrhein-Westfalen. All animal care and protocols adhered to national (Tierschutzgesetz) and European (Directive 2010/63/EU) laws and regulations as well as European Federation of Animal Science Associations (FELASA) http://www.felasa.eu/. Animals were euthanized using the cervical dislocation method. *Ptf1a^wt/Cre^;Kras^wt/LSL-G12D^;p53^fl/fl^* (*CKP*) mice have been described previously (24). Mice were on a mixed C57BL/6;129/Sv background. FKPC2GP is generated by crossing *Ptf1a^wt/Flp^;Kras^wt/FSF-G12D^;p53^frt/frt^* (FKP) mice to *Gt(ROSA)26Sor^tm3(CAG-Cre/ERT2)Das^* (*R26^FSF-CAG−CreERT2^*) and *Gt(ROSA)26Sor^tmloxP-STOP-loxP-GP-IRES-YFP^* (*R26^LSL-GP^*) (25) strains.

All animals were numbered, genotypes were revealed and animals then assigned to groups for analysis. For treatment experiments mice were randomized. None of the mice with the appropriate genotype were excluded from this study. Details of original and interbred mouse strains are listed in **Table S4**.

### Cell culture and treatments

Human PDAC cell lines, PaTu8988T and MiaPaCa2 were purchased from the American Type Culture Collection. Stable cell lines for *GRN* suppression were established by transfecting *GRN* shRNA into Patu8988T and MiaPaCa2. Scramble shRNA was included as negative control (nc) for transfection. Please refer to the **Supplementary Experimental Procedures** for additional detail.

For GP82, it was established from the tumor of a FKPC2GP mouse no. 82. After enzymatic digestion of the tumor, the desegregated cells were grown and maintained in 10% FBS-supplemented DMEM. After third passage, cell expression of epithelial marker EpCAM and fibroblast marker α-SMA were assessed by immunofluorescence staining and flow cytometry, and confirmed an enrichment of epithelial cells with <1% contamination of fibroblasts.

### IF staining and flow cytometric analysis

For intracellular PGRN expression, cells were permeabilized with ice-cold 0.1% saponin and then incubated with mouse anti-human PGRN(17, 26), or equal amount of corresponding isotype control, following by FITC-goat anti-mouse antibody (BD biosciences). CD3, CD8, CD45.1 on T cells, cells were stained with corresponding antibodies or equal amount of corresponding isotype controls. For intracellular granzyme B, TNF-a and IFN-g in T cells, or LCMV-gp33 in tumor cells, cells were fixed with 4% paraformaldehyde for 10 min at 37°C. After washing twice with PBS, cells were permeabilized with 0.1% Saponin for 20 min and then stained with antibodies and corresponding isotype. Details of primary antibodies are listed in **Table S5**. Cells were then washed, resuspended, and subjected to analysis. Expression of corresponding molecules of 10,000 viable cells was analyzed by flow cytometry (FACSCelasta; BD Biosciences) as mean fluorescence intensity (MFI). Raw data were analyzed using FlowJo software version 7.5.5 (Tree Star Inc., Ashland, OR).

### Enzyme-linked immunosorbent assay

Soluble PGRN levels in human plasma samples and culture supernatants were detected by a human PGRN ELISA kit (Adipogen Inc.). Please refer to the **Supplementary Experimental Procedures** for additional detail.

### *In vivo* antibody treatment on *CKP*

*CKP* mice were injected with or without mouse immunoglobulin (mIg) or anti-PGRN antibody (clone A23, (27)) twice weekly at 50 mg/kg in PBS, via intraperitoneal injection at four weeks of age for two consecutive weeks. Mice were weighted twice weekly throughout experiment. Mice with >15% body weight loss were terminated even the endpoint was not reached. At the endpoint, serum, tumors and organs were collected and processed for histological and immunohistochemical analysis. Tumor burden was measured by establishing gross wet weight of the pancreas/tumor.

### Real-Time quantitative Reverse-Transcription Polymerase Chain Reaction

Real-time quantitative PCR (qPCR) was performed by Roche LightCycler® 480 using LightCycler® 480 SYBR Green I Master Kit (Roche GmbH, Germany). Please refer to the **Supplementary Experimental Procedures** for additional detail.

### Immunofluorescence staining

Please refer to **Table S6** for primary antibody list and the **Supplementary Experimental Procedures** for additional detail.

### Lysosomal staining

Lysosomal staining was performed using Cytopainter LysoGreen indicator reagent (Abcam) according to manufacturer’s instructions. Please refer to the **Supplementary Experimental Procedures** for additional detail.

### Isolation of T cells from P14-TCR-Tg mice

Spleens harvested from P14-TCR-Tg mice were homogenized, and then lysed by Ammonium-Chloride-Potassium lysis buffer. T cells were negatively selected by MACS (Miltenyi Biotec) according to manufacturer’s instructions. After isolation, T cells were labeled with 5uM CFSE (Thermofisher), and resuspended 10% FBS-supplemented RPMI medium in pre-stimulated with 20ng/ml IL-2 (Peprotech) for 1hr, before co-culture with GP82 cell line or orthotopic injection in C57BL/6J mice.

### Co-culture of LCMV-gp33-reactive T cells and GP82 cells

GP82 cells were treated with or without tamoxifen to induce LCMV-gp33 expression for at least 5 days, and then treated with or without PGRN Ab or mIg to suppress PGRN levels. After 2-day treatment, cells were harvested and seed to 6-well with 10% FBS-supplemented DMEM and were ready to co-culture. Then, LCMV-gp33-reactive T cells were isolated, and then cultured with or without GP82 cell lines at ratio of 8:1, in 10% FBS-supplemented DMEM for 2 days. Cells were then photographed under microscope, and then harvested for anti-tumor cytotoxicity by propidium iodide (PI) staining. Besides, T cell activity was assessed by staining for CD8, granzyme B, TNF-a, IFN-g by flow cytometry.

### *In vivo* antibody treatment on orthotopic model of GP82 cells in C57BL/6J mice

GP82, primary cell line derived from one of the FKPC2GP tumor, was transplanted orthotopically into the pancreas of C57BL/6J mice with needle injection under ultrasound imaging guidance. After orthotopic transplantation of GP82 cells, ultrasound imaging was performed once a week to monitor tumor growth. Once tumor volume reached 100mm^3^, intraperitoneal injection of tamoxifen (75mg/kg) will be performed to induce LCMV-gp33 expression. After second injection of tamoxifen (2 days after first injection), mouse immunoglobulin (mIg) or anti-PGRN antibody (clone A23, (27)) twice weekly at 50 mg/kg in PBS, via intraperitoneal injection for 2 weeks. After the second treatment of A23 or mIg, LCMV-gp33-reactive T cells isolated from P14-TCR-Tg mice were injected intravenously. Mice were weighted twice weekly throughout experiment, and tumor growth was monitored by ultrasound imaging once a week. Mice with >15% body weight loss were terminated even the endpoint was not reached. At the endpoint, serum, tumors and organs were collected and processed for immunohistochemical and flow cytometric analysis.

### Statistical data analysis

All statistics were performed using GraphPad Prism 6.0 (GraphPad Software, La Jolla, CA). Survival data was analyzed by Log-rank (Mantel-Cox) test, while correlation analysis was by Spearman’s rank correlation coefficient. Chi-square test was used to assess the independency between two categorical variables. Two-tailed Mann-Whitney test was applied for non-normally distributed data comparison between two groups. For multiple group comparison, Kruskal-Wallis test was used. Data are represented as mean ± S.D. * p < 0.05; ** p < 0.01; *** p < 0.001

## RESULTS

### PGRN exerts distinct functions on tumor cells and macrophages in human PDAC

To understand the role of PGRN in human PDAC development, we delineated its expression pattern at different stages in PDAC specimens (Essen cohort, n=53). PGRN expression was observed in preneoplastic cells and remained throughout the malignant transformation **(Figure 1a)**. The staining was quantified and the number of PGRN+ cells was significantly higher in tumor than non-tumor areas **(Figure S2a)**. Patients were dichotomized into high and low PGRN expression groups (**Figure 1b)** with median of PGRN+ cells as cutoff. Low PGRN expression showed significantly superior survival than the high expression group (Median overall survival: high vs low PGRN expression group: 9 vs 21mo, **Figure S2b**). We validated the association of PGRN expression with survival in an independent cohort from Nijmegen (n=31,**Figure S2c, d)**.

**Figure 1.**
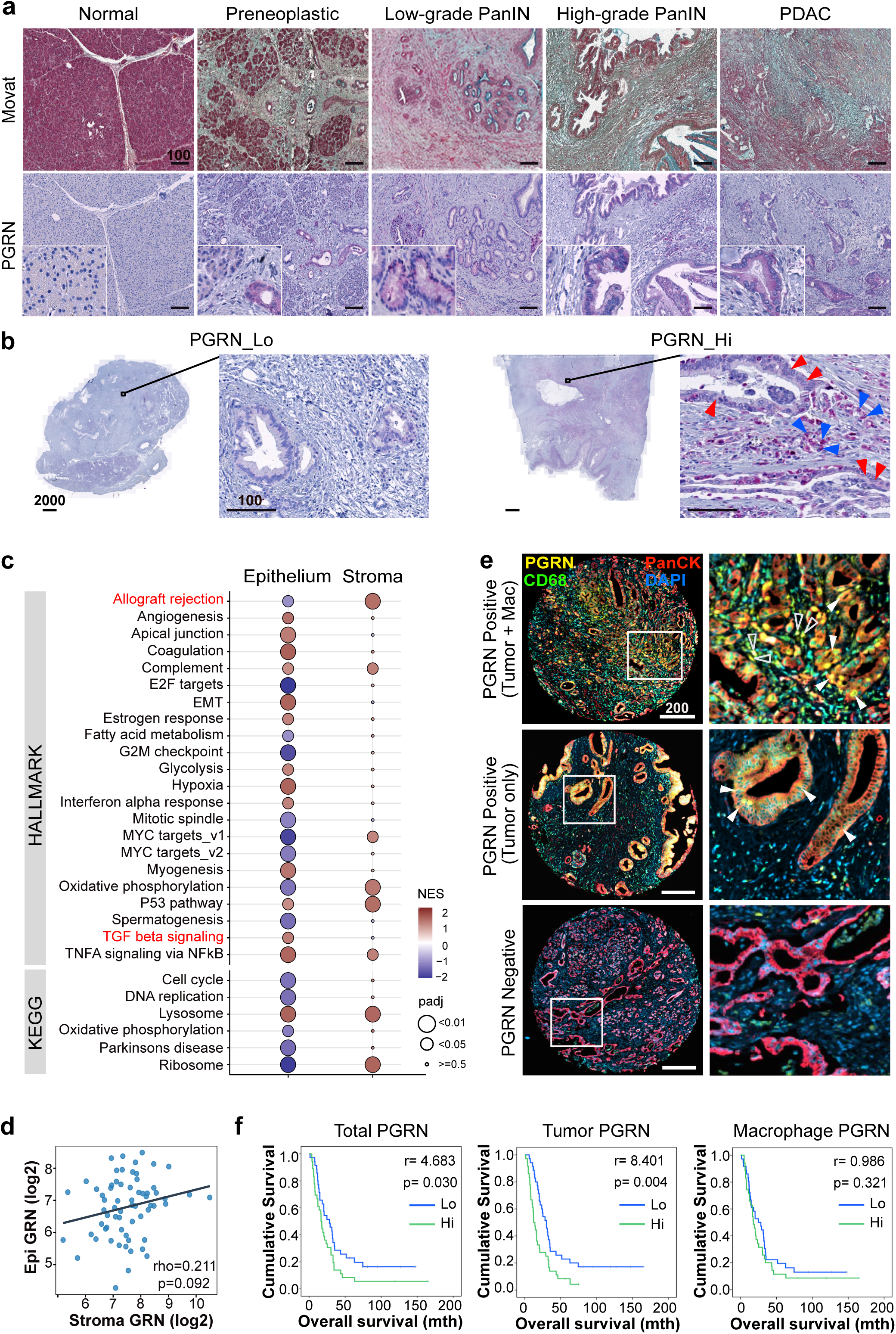
PGRN exerts different functions on tumor cells and macrophages in human PDAC. **(a,b)** Essen cohort (n=53). **(a)** PGRN expression pattern during human PDAC development. Movat and IHC staining of PGRN in normal pancreas, preneoplastic low- and high-grade PanIN stages, and tumorous tissues of PDAC patients. **(b)** IHC staining of representative patient specimen with high and low PGRN levels as defined by the quantification of Definiens, with median of number of PGRN+ cells as cutoff. **Right panel:** PGRN expression in both tumor and stromal compartments of PDAC. Red arrowheads indicate PGRN+ tumor cells; blue arrowheads indicate PGRN+ stromal cells. **(c, d)** Maurer *et al* dataset (GSE93326, n=65). **(c)** GSEA result of hallmark and KEGG pathways in PDAC epithelium and stroma samples reveals different enriched gene sets for high *GRN* in epithelium and stroma. Enrichment was tested against the differential expression profile of *GRN*-high (n=32) versus *GRN*-low (n=32) in epithelium and stroma samples separately. Pathways with padj < 0.01 from either epithelium or stroma groups were shown with their NES and padj values. A complete list of significant pathways is shown in **Table S7**. Gene sets of interest are highlighted in red. **(d)** No significant correlation between epithelium (tumor) and stroma *GRN* expression levels (Log2CPM) in 65 pairs of PDAC specimens. **(e, f)** CONKO-001 cohort (n=71). Tissue microarrays were stained for PGRN, CD68 (macrophage), PanCK (tumor) and DAPI by multiplex immunofluorescent (mIF) staining and quantified by Definiens. **(e)** Representative tissue cores showing PGRN expression in: both tumor cells (Tumor) and macrophages (Mac), in tumor cells only, and negative in both cell types. **Right column:** Filled arrowheads indicate PGRN+ tumor cells; Hollow arrowheads indicate PGRN+ macrophages. **(f)** PDAC samples were categorized into high (n=35) and low (n=36) expression groups (cutoff: median of the number of PGRN+ cells) based on the PGRN expression in all the cells (Total PGRN), tumor cells (Tumor PGRN, PGRN+PanCK+) only, and macrophages (Macrophage PGRN, PGRN+CD68+) only. Kaplan-Meier overall survival plots according to PGRN expression level in different cell compartments. Scale bar unit: µm

Intriguingly, PGRN expression was observed in both tumor and stromal compartments in both Essen **(Figure 1b, right panel)** and Nijmegen cohorts **(Figure S2d)**. To dissect the roles of PGRN in tumor and stroma separately, we analyzed the RNA-sequencing dataset of Maurer *et al*(19) (GSE93326, n=65), where PDAC malignant epithelium and stroma were procured by laser capture microdissection. Cell type deconvolution was performed with the transcriptomes derived from high *GRN* (*n*=32) and low *GRN* (*n*=32) stroma samples using 43 pre-defined different cell types, showing that macrophages were enriched in high *GRN* stroma **(Figure S2e, Table S1)**. Macrophages also represent the cell type with the highest degree of variable importance in predicting high *GRN* stroma using random forest analysis **(Figure S2f)**.

Gene set enrichment analysis (GSEA) of the hallmark and KEGG pathways was performed and revealed that the pathways enriched for high *GRN* were different for epithelium and stroma **(Figure 1c**, **Table S7)**. Notably, in epithelium but not stroma, high *GRN* showed down-regulation in the allograft rejection gene set, which implies a role in immune evasion; as well as an enrichment in TGF-β signaling, which contributes to immune exclusion and evasion in various cancer types(28). Besides, no significant correlation was observed between *GRN* expression in epithelium and stroma **(Figure 1d)**.

We performed multiplex immunofluorescence (mIF) to distinguish PGRN expression derived from tumor cells (PanCK+) and macrophages (CD68+) in tissue microarrays generated from the adjuvant clinical phase III CONKO-001 cohort (gemcitabine vs no treatment, here: trial arm without chemotherapy, n=71)(29, 30). Three different patterns of PGRN expression were observed: PGRN positive signals in 1) both tumor and macrophages; 2) in tumor only; and 3) negative in both cell compartments **(Figure 1e)**. Survival analysis showed that high PGRN expression in total cells and tumor cells both predicted poor survival, while PGRN expression in macrophages did not predict survival **(Figure 1f)**. Besides, we analyzed the association of tumor PGRN with various clinicopathological parameters and immune markers that were characterized previously in this cohort(29, 30). Interestingly, high tumor PGRN level was significantly associated with CD8+ cell abundance in negative manner **(Table S8)**, implying a potential immunoregulatory role of PGRN expression in primary PDAC.

### PGRN+ tumors of PDAC patients exhibit lower levels of MHCI expression and CD8 infiltration

To address the spatial interaction between tumor PGRN and CD8+ cells, we performed multiplex immunofluorescence (mIF) in 8 cases of human PDAC. Intratumoral heterogeneity was observed in all specimens with high and low PGRN-expressing tumor areas **(Figure 2a)**. Notably, in high PGRN-expressing tumor regions we rarely detected CD8+ cells, while the opposite was found in low PGRN-expressing tumor regions of the same tumors **(Figure 2b)**. Next we determined if tumor PGRN expression is associated with cytotoxic activity of CD8+ cells, by including cytotoxic marker granzyme B (GzmB) and MHCI molecule HLA-A in the mIF. In low PGRN-expressing tumor region, PGRN−/PanCK+ tumor cells showed stronger expression of MHCI molecule HLA-A. Also, higher CD8 infiltration could be observed, in which a significant portion of them was GzmB+ **(Figure 2c)**. On the contrary, PGRN+/PanCK+ tumor cells expressed no, or only low levels of MHCI, and infiltrating CD8+ and GzmB+ cells were scarce **(Figure 2c)**. Spatial analysis showed that percentage of MHCI+ cells in PGRN−/PanCK+ tumor cells was significantly higher than the PGRN+ counterparts. Besides, significantly more CD8+/GzmB+ cells could be found in close proximity (≤50µM radial distance) of PGRN−/PanCK+ cells **(Figure 2d)**, implying a potential regulatory role of PGRN in tumor immunogenicity and anti-tumor cytotoxicity in PDAC.

**Figure 2.**
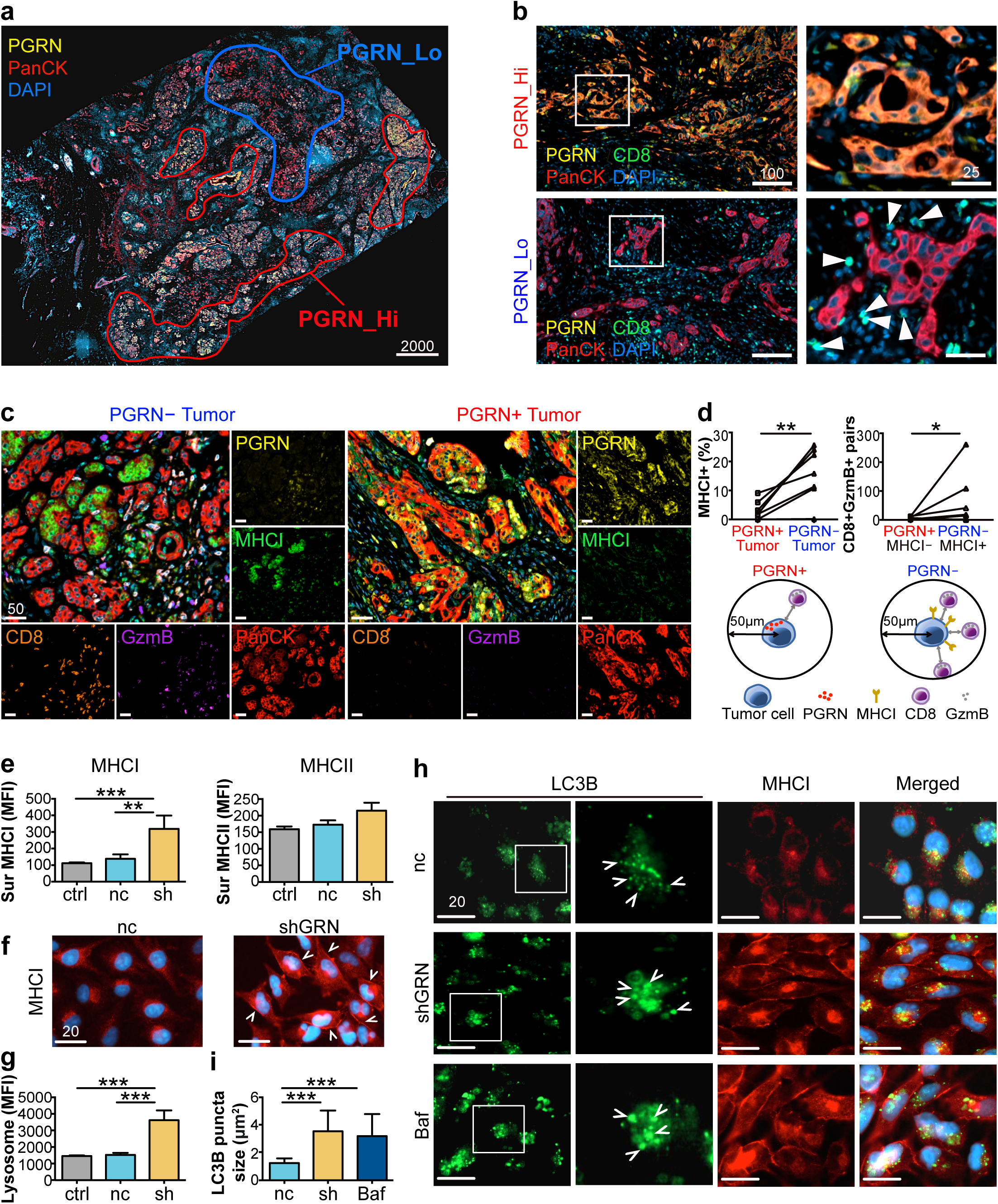
PGRN regulates MHCI expression via autophagy in human PDAC. (**a**) Representative mIF image demonstrates the intratumoral heterogeneity of PGRN expression in human PDAC, where high and low PGRN-expressing tumor regions were observed in the same specimen. **(b**) mIF staining of PGRN (yellow), CD8 (green) and PanCK (red) in human PDAC shows increased CD8 infiltration in low PGRN-expressing tumor area. **(c)** Differential MHCI (HLA-A) expression in PGRN+ and PGRN- tumors, and the infiltration of GzmB+CD8+ cells in their corresponding neighborhoods. **(d)** Automated computational analysis showing the percentage of MHCI+ cells in PGRN+/PanCK+ and PGRN-PanCK+ populations, and the number of CD8+GzmB+ cells in proximity (<50µM radical distance) of PGRN+/MHCI-/PanCK+ or PGRN-/MHCI+/PanCK+ tumor cells in human PDAC (n=8). **(e**) Surface MHCI (HLA-A/B/C) and MHCII (HLA-DR) expression on human PDAC cell line MiaPaCa2 upon *GRN* suppression is assessed by flow cytometry (n=4). **(f)** IF staining of MHCI marker HLA-A/B/C reveals augmented surface MHCI expression on MiaPaCa2 upon *GRN* suppression. White arrowheads indicate the membraneous staining of MHCI. **(g)** Lysosome content in MiaPaCa2 upon *GRN* suppression is assessed by staining with fluorescent LysoGreen Indicator and measured by flow cytometry (n=4). **(h, i)** IF staining of MHCI (red) and LC3B (green) in MiaPaCa2 cells upon GRN suppression or treated with autophagy inhibitor Bafinomycin (Baf, 100nM, 24h). **(i)** Average size of LC3B puncta of 30 cells of each treatment was measured by ZEN software. ctrl: parental PDAC cells; nc: shRNA scrambled control; sh/shGRN: *GRN* shRNA; Baf: Bafinomycin. MFI: mean fluorescence intensity. **p<0.01, **p<0.01, ***p<0.001. Scale bar unit: µm

### PGRN suppression leads to surface MHCI upregulation and dysfunctional autophagy in human PDAC cells

To assess the cell-autonomous effect of PGRN on MHCI expression, we examined the surface MHCI molecules HLA-A/B/C, on human PDAC cell lines upon *GRN* suppression using RNA interference. We selected two human PDAC cell lines, MiaPaCa2 and PaTuT cells, with relatively high PGRN expression levels **(Figure S3a-c)**. Surface expression of MHCI on both cell lines was significantly augmented upon RNAi-mediated knockdown of *GRN* **(Figure 2e, S3d).** MHCII was also examined and there was no significant change in surface MHCII expression upon *GRN* suppression **(Figure 2e, S3d)**, confirming that the PGRN effect was restricted to MHCI. By IF, membranous staining of MHCI was observed clearly in *GRN* suppressed cells, but not in the controls **(Figure 2f, S3e).**

It was demonstrated that pancreatic cancer cells down-regulated surface MHCI by autophagy-mediated degradation(31). Coincidentally, PGRN was shown to regulate autophagy in various cell types(32, 33). PGRN-deficient mice showed defective degradation of autophagosomes under starvation(34), which was mediated by lysosomal enzymes cathepsin D(35). Notably, our GSEA data revealed in high *GRN* tumor an enrichment of lysosome pathway **(Figure 1c,Table S7c)** with multiple cathepsin (CTS) genes including CTSD as the leading edge genes. Therefore, we reasoned that PGRN might control surface expression of MHCI through autophagy. We measured lysosome marker LAMP1 in the *GRN*-suppressed cell lines after nutrient starvation (1% FBS). Accumulation of lysosomes was significantly increased in sh*GRN* transfectants **(Figure 2g, S3f)**. Next, we co-stained MHCI and autophagosome marker LC3B in the cells under nutrient-starved condition. Upon *GRN* suppression, the number and size of LC3B puncta increased significantly when compared to the controls**(Figure 2h,i,S3g,h)**. The increase in LC3B punctae size was comparable to the cells treated with the V-ATPase inhibitor bafilomycin A1, which blocks the degradation of autophagosomes**(Figure 2h,i,S3g,h)**, suggesting that the LC3B accumulation observed in *GRN* suppression is caused by impaired clearance of autophagosomes. Besides, in control cells, MHCI and LC3B clearly co-localized (**Figure 2h,S3g)**, supporting the previous findings that PDAC cells down-regulated surface MHCI molecules by autophagy-dependent lysosomal degradation(31). These findings suggest that PGRN suppression in PDAC cells restores cell surface MHCI expression by inhibition of autophagy-mediated degradation.

### PGRN blockade suppresses tumorigenesis *in vivo*

Next, we investigated the effect of PGRN inhibition on PDAC development *in vivo* using an antibody-based PGRN blockade strategy. Anti-PGRN antibody (Ab) was previously shown to neutralize soluble PGRN (sPGRN) in liver cancer(16,27,36). Since PGRN is a secreted glycoprotein regulating its own expression in autocrine/paracrine manner, neutralization of sPGRN reduces both extracellular and intracellular PGRN(27). We treated MiaPaCa2 and PaTuT cells with anti-PGRN Ab *in vitro*, and both cellular and secretory PGRN levels were significantly reduced upon Ab treatment **(Figure S4)**, supporting subsequent targeting of cellular and circulating PGRN *in vivo*.

Since PGRN expression was observed in preneoplastic lesions in human PDAC, we hypothesized a functional role in early PDAC development. We delineated PGRN expression pattern during PDAC development in a well-characterized spontaneous endogenous PDAC mouse model (termed *CKP*) with conditional oncogenic *Kras^G12D^* mutation and loss of *Tp53*(24, 37), which allows longitudinal characterization of tumor evolution. Pancreata were harvested at the stages preneoplasia (4 weeks), early (6 weeks) and advanced PDAC (10 weeks). IHC revealed PGRN expression in preneoplastic lesions recapitulating expression patterns in human PDAC. As lesions progressed to early PDAC, PGRN levels reached the maximum, and slightly decreased in advanced stage **(Figure 3a)**.

**Figure 3.**
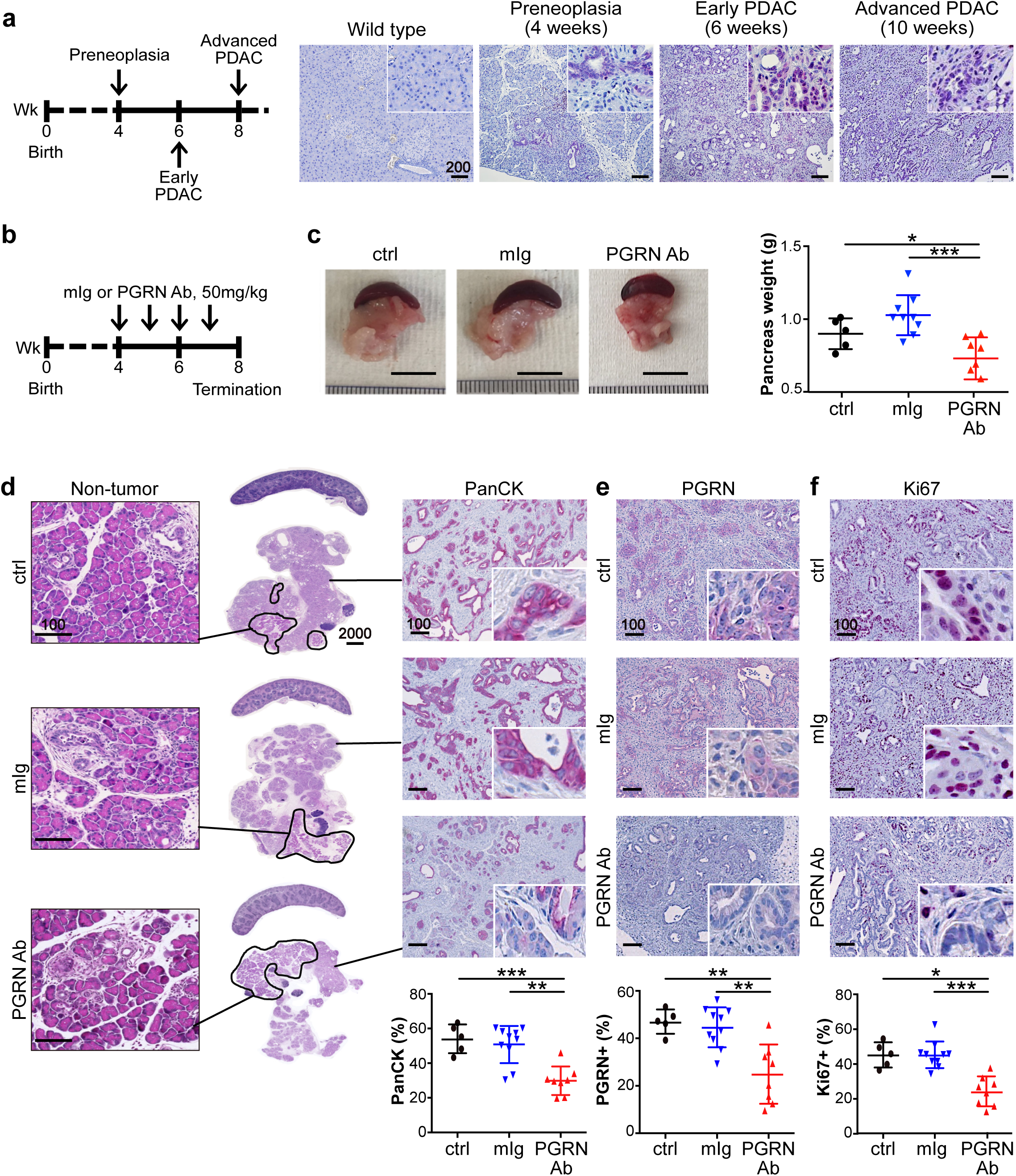
*In vivo* PGRN blockade suppresses tumor initiation and progression in mouse PDAC. (**a**) IHC staining of PGRN in pancreas of wild type mice and *CKP* mice with preneoplasia (4 weeks), early (6 weeks) and advanced (10 weeks) PDAC. **(b)** Timeline for treatment of *CKP* mice with mouse isotype (mIg) or anti-PGRN antibody (PGRN Ab), 50mg/kg. **(c)** Representative pictures of tumors and spleens from mIg-treated (n=10), PGRN Ab-treated (n=8), and untreated (CTL, n=5) *CKP* mice. Right panel: Weight of pancreas of *CKP* mice upon dissection. **(d)** PGRN blockade suppresses tumorigenesis in *CKP* mice. Middle panel: H&E staining of pancreas and spleens of *CKP* mice. Non-tumorous pancreatic tissues (left panel) are highlighted by black lines. Right panel: IHC staining of panCK in the *CKP* pancreata. The lower panel shows the percentage of panCK+ cells in the whole pancreata. IHC staining of **(e)** PGRN and **(f)** Ki67 in *CKP* tumors. The lower panels show the percentage of PGRN+ and Ki67+ cells in the whole pancreas. Mean ± SD is shown. *p<0.05,**p<0.01,***p<0001. Scale bar unit: µm

Anti-PGRN Ab treatment started when mice were 4 weeks old (preneoplasia) and was terminated at 6 weeks, when PDAC formation is frequently observed in this model **(Figure 3b)**. Anti-PGRN Ab treatment significantly alleviated tumor burden as compared to controls in terms of tumor weight **(Figure 3c)** and proportion of tumorous (PanCK+) tissues **(Figure 3d)**. IHC analysis confirmed that PGRN+ cells in Ab-treated tumors were significantly reduced **(Figure 3e)**. Notably, Ki67+ tumor cells were also significantly reduced, indicating reduced tumor proliferation **(Figure 3f)**. Thus, PGRN blockade prominently halts tumor initiation and progression in this aggressive genetic model of spontaneous PDAC development.

### *In vivo* PGRN blockade revives CD8 anti-tumor cytotoxicity

Next, we focused on the effect of PGRN blockade on tumor immune microenvironment. We found the number of infiltrating CD3+ and CD8+, but not CD4+ cells, significantly increased upon Ab treatment **(Figure 4a)**. The increase of CD8+ cells was not solely due to the delayed tumor progression. We previously delineated the dynamic of immune landscape throughout PDAC development in this model(37) and found that CD8+ cells contributed to less than 4% of cells in tumors (re-quantified in whole-tissue scale, **Figure S5a**). Upon Ab treatment, we observed up to 10% CD8+ cells in the tumors **(Figure 4a)**, indicating that PGRN blockade promotes additional CD8+ cell infiltration.

**Figure 4.**
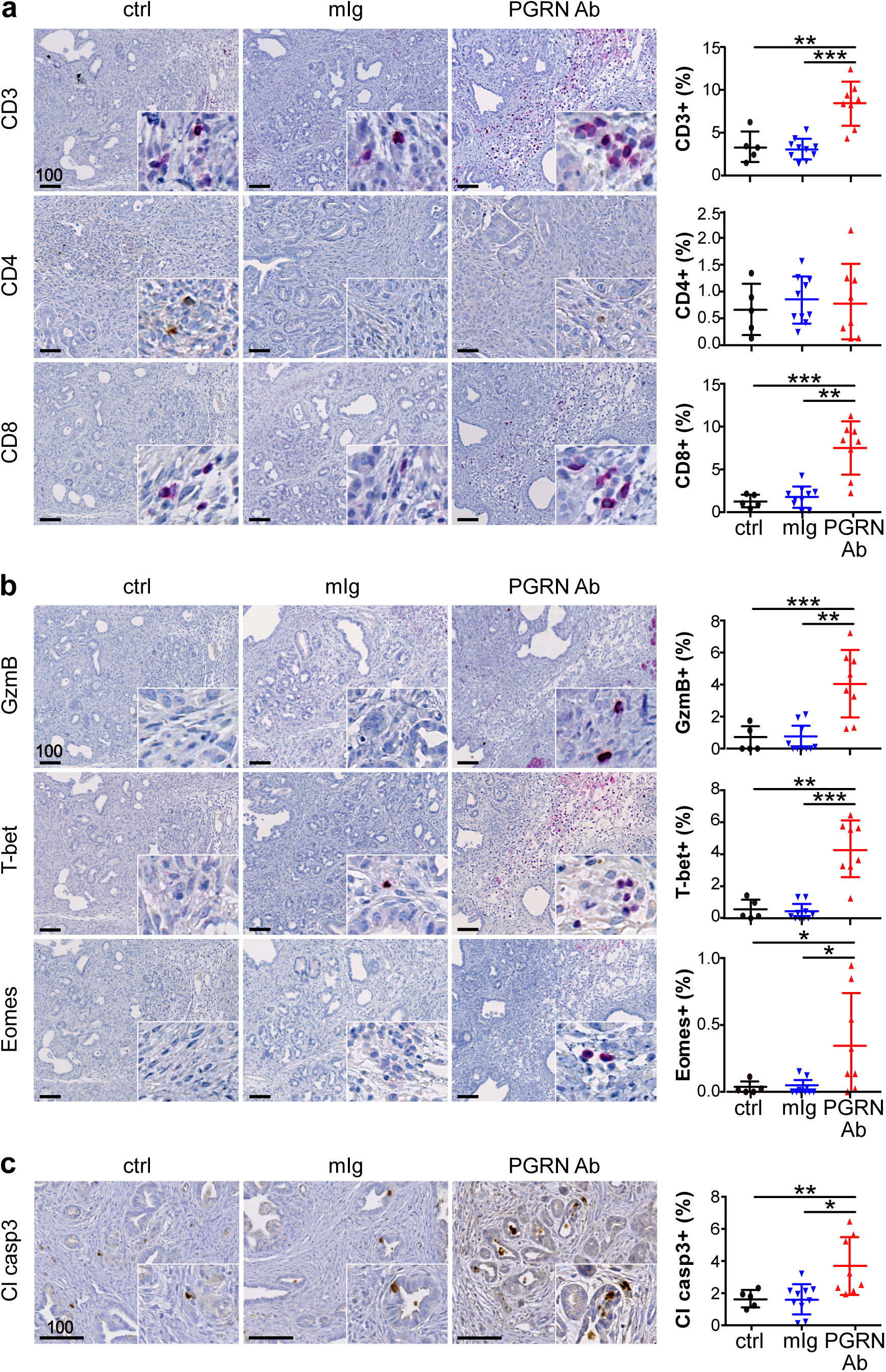
*In vivo* PGRN blockade revives CD8 anti-tumor cytotoxicity. IHC staining of **(a)** T cell markers CD3, CD4 and CD8; **(b)** cytotoxic markers granzyme B (GzmB), T-bet and Eomes; and **(c)** apoptotic marker cleaved caspase 3 (Cl casp3) in *CKP* tumors treated with or without PGRN Ab or mIg. ctrl: n=5; mIg: n=10; PGRN Ab: n=8. The right panels show the percentages of positive cells in the whole tumors. *p<0.05,**p<0.01,***p<0001. Scale bar unit: µm

Importantly, the expression of cytotoxic markers granzyme B, T-bet and Eomes were also significantly augmented**(Figure 4b)**, implying that PGRN Ab enhanced the cytotoxicity of the infiltrating CD8+ T cells. The anti-tumor cytotoxicity induced by PGRN blockade was confirmed by increased level of cleaved caspase-3 in tumor cells **(Figure 4c)**.

Since TGF-β signaling was upregulated in high *GRN* epithelium, we assessed the effect of PGRN blockade on regulatory T cells. The number of foxp3+ cells was significantly suppressed upon PGRN blockade **(Figure S5b)**. However, since the foxp3 expression level was low in the *CKP* model, we consider it unlikely that the revived CD8 cytotoxicity was caused by the suppression of regulatory T cells.

### *In vivo* PGRN blockade suppresses M2 polarization but not fibroblast accumulation

It was reported that macrophage-derived PGRN associates with M2 phenotypes, and most importantly blocks the infiltration of CD8+ cells in metastatic PDAC by promoting fibrosis (8, 9). Here, we examined the effect of PGRN blockade on TAM infiltration and fibrosis. Pan-macrophage marker F4/80 level was not significantly changed upon PGRN blockade **(Figure S5c)**. However, TAM marker MRC1 level was greatly reduced, while M1 markers phospho-STAT1 and iNOS increased upon treatment **(Figure S5c)**, indicating that PGRN blockade skewed macrophage polarization from M2 to M1 phenotype. However, CD4 infiltration remained unchanged upon PGRN blockade **(Figure 4a)**, indicating that the increase in M1 macrophages might not to be sufficient to attract the CD4+ T cells.

Our results regarding the effect of PGRN on M2 polarization echo the previous findings by Nielsen *et al* in metastatic PDAC, in which macrophage-derived PGRN induced fibrosis in non-cell autonomous manner and blocked CD8 infiltration(9). Therefore, we also examined the abundance of fibroblasts in the PGRN Ab-treated tumors. Paradoxically, no significant difference was observed in fibroblast accumulation (α-SMA+ cells) in the tumors **(Figure S5d)**, suggesting that the revived CD8 infiltration and activation induced by PGRN blockade is not mediated by fibrosis reduction.

### *In vivo* PGRN blockade restores tumor MHCI expression that is spatially associated with increased CD8 cell infiltration

Next, we assessed the expression of MHCI molecule H-2Db in the *CKP* tumors. H-2Db expression, which appeared low or absent in tumor cells, was significantly restored upon PGRN Ab treatment **(Figure 5a)**. However, MHCII expression was not changed upon PGRN treatment **(Figure 5a)**, which echoes our earlier findings on the unchanged CD4 infiltration. We also examined lysosome marker LAMP1 and autophagosome marker LC3B in PGRN Ab-treated *CKP* tumors. Both markers were significantly increased upon PGRN blockade **(Figure S6a)**, implying the presence of dysfunctional autophagy that might be involved in the PGRN blockade-mediated MHCI expression.

**Figure 5.**
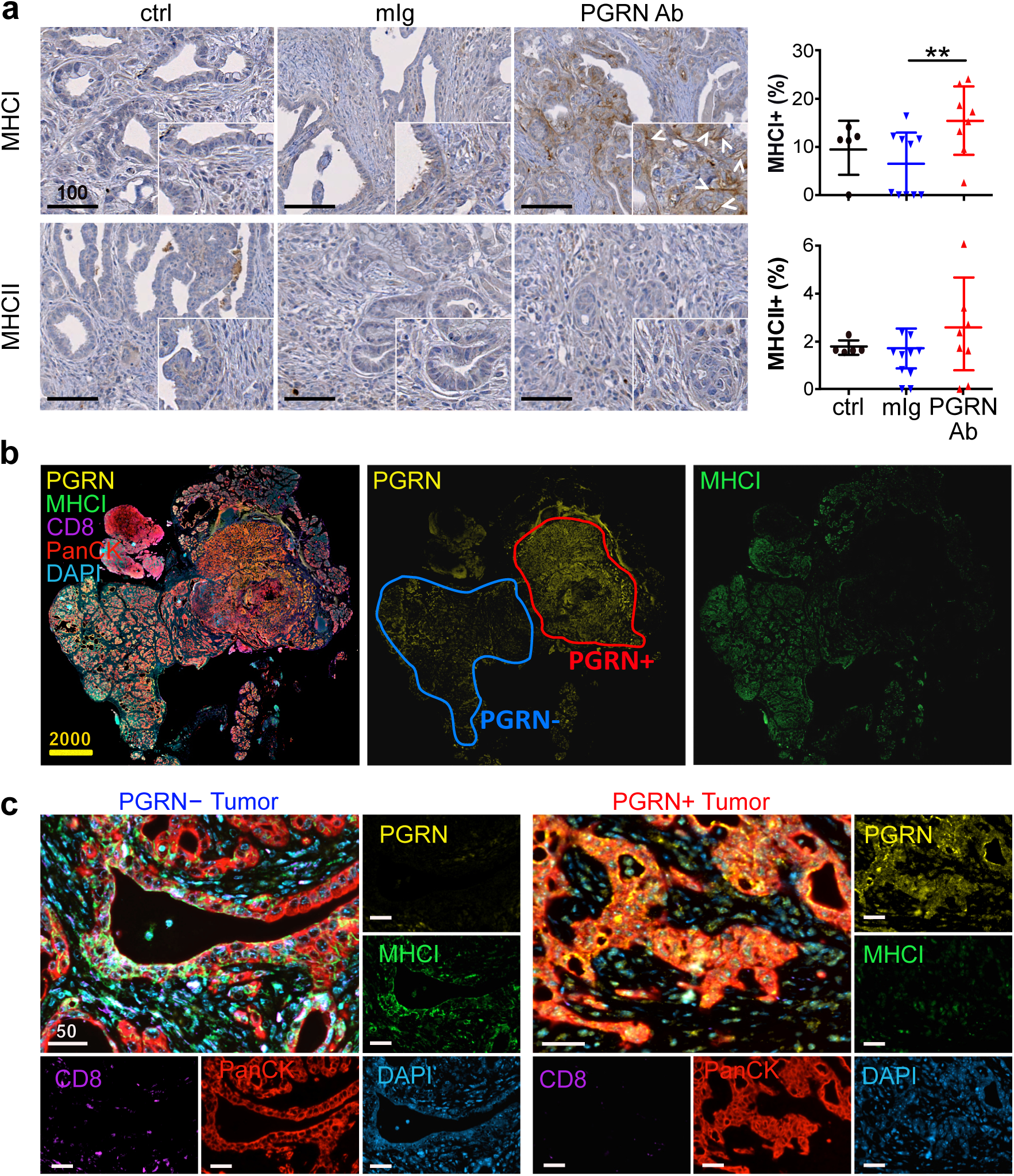
*In vivo* PGRN blockade restores tumor MHCI expression that is spatially associated with increased CD8 cells. **(a)** IHC of MHCI marker H-2Db and MHCII in *CKP* tumors treated with or without PGRN Ab or mIg. ctrl: n=5; mIg: n=10; PGRN Ab: n=8. The right panels show the percentages of positive cells in the whole tumors. **(b)** mIF of PGRN (yellow), MHCI (green), CD8 (purple) and PanCK (red) in PGRN Ab-treated *CKP* tumors (n=8). Intratumoral heterogeneity of PGRN expression was observed, in which PGRN+ and PGRN- regions were depicted in the representative tumor. PGRN and MHCI show opposite staining pattern in the PGRN Ab-treated tumor. **(c)** mIF showing the differential MHCI and CD8 expression in PGRN+ and PGRN- tumor regions. **p<0.01. Scale bar unit: µm

mIF was performed to illustrate the spatial association among PGRN, MHCI and CD8 cells. We observed heterogeneous expression patterns of PGRN and MHCI across the PGRN Ab-treated tumor (n=8, **Figure 5b**). Notably, MHCI expression was absent in the PGRN+ region with advanced PDAC tumor, while it was observed in the PGRN− region with mostly preneoplastic lesions **(Figure 5b)**. Spatial analysis demonstrated that the proportion of MHCI+ cells was significantly higher in PGRN− tumor cells **(Figure S6b)**. Besides, significantly more CD8+ cells were found in the vicinity (≤50µm radical distance) of PGRN−MHCI+ tumor cells when compared to PGRN+ counterparts **(Figure 5c, S6b)**.

### *In vitro* PGRN blockade promotes CD8 anti-tumor cytotoxicity via MHCI regulation

Next, we investigated if PGRN-induced MHCI restoration could effectively lead to functional effects on anti-tumor cytotoxicity. Due to low mutational burden, neoantigen levels in PDAC are relatively low. Thus, we generated a dual recombination (cre/lox;flp/frt) next-generation mouse model with spatial and temporal controlled antigen expression, namely gp33 of LCMV, in tumor cells. Here, flp-mediated activation of KrasG12D and loss of Tp53 in pancreatic progenitors (*FKP* model) leads to spontaneous PDAC development mirroring tumorigenesis in the cre-mediated *CKP* model. We interbred *FKP* mice with Rosa26-LSL-GP and Rosa26-FSF-CreERT2 mice (*FKPC2GP*), in which CreERT2 expressing pancreatic cells express LCMV-gp33 upon tamoxifen administration **(Figure 6a)**. We established a primary pancreatic cancer cell line, GP82, from a *FKPC2GP* mouse and induced LCMV-gp33 expression in GP82 *in vitro*. The induced GP82 cells were co-cultured with LCMV-gp33-reactive T cells isolated from the spleen of P14-TCR-Tg mice **(Figure 6b)**, to assess the effect of PGRN Ab on tumor-specific cytotoxicity.

**Figure 6.**
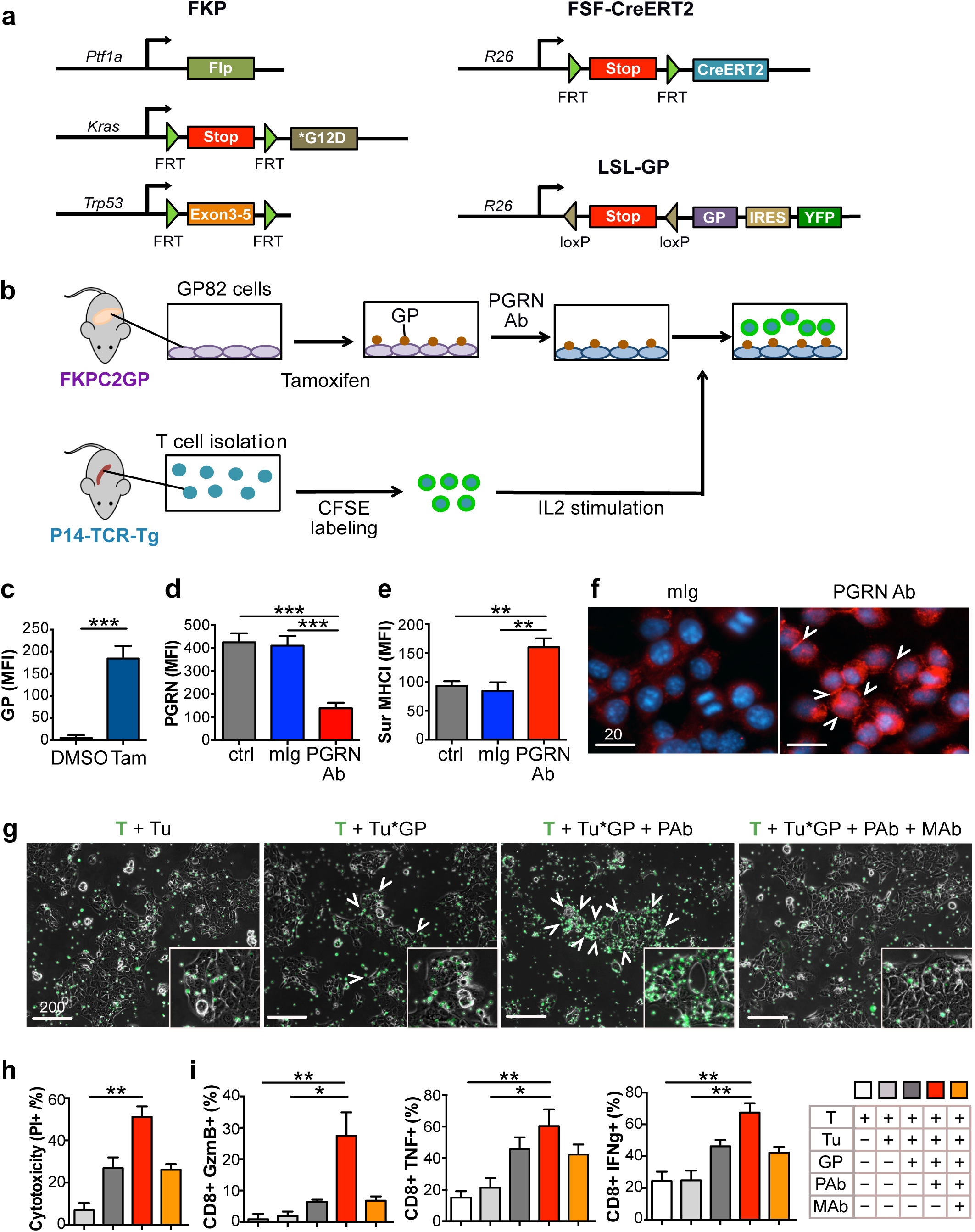
*In vitro* PGRN blockade promotes CD8 anti-tumor cytotoxicity via MHCI regulation. Genetic strategy to induce spatially and temporally controlled GP33 expression by tamoxifen-mediated activation of CreERT2 in pancreatic cells harboring mutant *Kras^G12D^* and loss of *Tp53*. *Ptf1a^wt/flp^;Kras^wt/FSF-G12D^;p53^frt/frt^* (FKP) mice crossed to *Gt(ROSA)26Sor^tm3(CAG-Cre/ERT2)Das^* (*R26^FSF- CAG−CreERT2^*) and *Gt(ROSA)26Sor^tmloxP-STOP-loxP-GP-IRES-YFP^* (*R26^LSL-GP^*) strains to generate FKPC2GP mice. Experimental setup for co-culture of GP82, LCMV-gp33-expressing cell line derived from FKPC2GP tumor, and the gp33-reactive T cells isolated from the spleen of P14-TCR-Tg mice. **(c**) LCMV-gp33 (GP) expression in GP82 cells treated with tamoxifen (25uM) or vehicle control DMSO for 2 days was assessed by flow cytometry (n=4). Tam: Tamoxifen. **(d**) Cellular PGRN level in GP82 cells treated with or without PGRN Ab or mIg (100ug/ml) was assessed by flow cytometry (n=4). **(e**) Surface expression of MHCI marker H-2Db on GP82 cells treated with or without PGRN Ab or mIg (100ug/ml) was assessed by flow cytometry (n=6). **(f**) IF staining of MHCI marker H-2Db of GP82 upon treatment with or without PGRN Ab or mIg (100ug/ml). White arrowheads indicate the membraneous staining of MHCI. **(g**) Microscopic images of GP82 cells and LCMV-gp33-reactive T cells (CFSE-labeled, green) after 2 days of co-culture. When anti-MHCI (H-2Db) neutralizing antibody (MHCI Ab) was included in the treatment, MHCI Ab was added 1h after PGRN Ab treatment. T cells were then added 1h after MHCI Ab treatment. White arrowheads indicate T cell clusters accumulated at GP82 cells. **(h**) Cytotoxicity level (PI+ %) of GP82 cells upon co-culture with LCMV-gp33-reactive T cells (n=6). Tu: LCMV-gp33-induced GP82 tumor cells; PAb: PGRN Ab (100ug/ml); MAb: MHCI Ab (100ug/ml). **(i**) Percentage of CD8+ cells that are positive for cytotoxic markers granzyme B (GzmB), TNF-a, and IFN-g (% in total T cells) assessed by flow cytometer (n=6). MFI: Mean Fluorescence Intensity. *p<0.05; **p<0.01; ***p<0.001. Scale bar unit: µm

We first confirmed LCMV-gp33 expression in GP82 cells upon tamoxifen treatment **(Figure 6c)** and treated them with PGRN Ab or mIg for 2 consecutive days. PGRN was significantly down-regulated upon Ab treatment **(Figure 6d)**. Congruent with our findings in *GRN*-suppressed human cell lines, the surface expression of MHCI molecule H-2Db on GP82 cells increased significantly by PGRN Ab treatment **(Figure 6e)**. IF staining also demonstrated that H-2Db expression in PGRN Ab-treated cells was greatly increased with membranous localization **(Figure 6f)**.

Upon co-culture, LCMV-gp33-reactive T cells were found to accumulate close to GP82 cells induced for LCMV-gp33 expression, but not to those without induction **(Figure 6g, S7a,b)**. Upon treatment with PGRN Ab, accumulation of T cells in the vicinity of tumor cells was markedly potentiated. However, the T cell accumulation was prominently abrogated upon MHCI blockade with H-2Db neutralizing antibody **(Figure 6g, S7a,b)**. Importantly, the cytotoxicity level of LCMV-gp33-induced cells upon PGRN Ab treatment was significantly increased as compared to control, and the PGRN Ab-induced cytotoxicity was abrogated upon addition of anti-MHCI neutralizing antibody **(Figure 6h, S7c)**. We next assessed T cell cytotoxic activity in the co-culture system. Quantities of granzyme B+, TNFa+, and IFNg+ CD8+ cells were all significantly increased in PGRN Ab treatment when compared to the controls **(Figure 6i, S7d)**, which again, was significantly diminished with the anti-MHCI neutralizing antibody, confirming the indispensable role of MHCI in the PGRN Ab-induced tumor-specific cytotoxic effect.

### *In vivo* PGRN blockade promotes antigen-specific T cell cytotoxicity against tumors

Next, we validated the above effect of PGRN Ab treatment *in vivo*. **Figure 7a** illustrates the experimental setup and treatment timeline. GP82 cells were transplanted orthotopically into C57BL/6J mice. Upon tumor formation confirmation by ultrasound imaging, tamoxifen was injected to induce LCMV-gp33 expression in the tumor. After two tamoxifen injections, PGRN Ab (n=4) or corresponding isotype control (mIg,n=4) was given twice a week for two weeks. LCMV-gp33-reactive T cells, freshly isolated from spleens of P14-TCR-Tg mice, were injected intravenously one day after the first administration of PGRN Ab or mIg. Mice were terminated one day after the last administration of Ab treatment.

**Figure 7.**
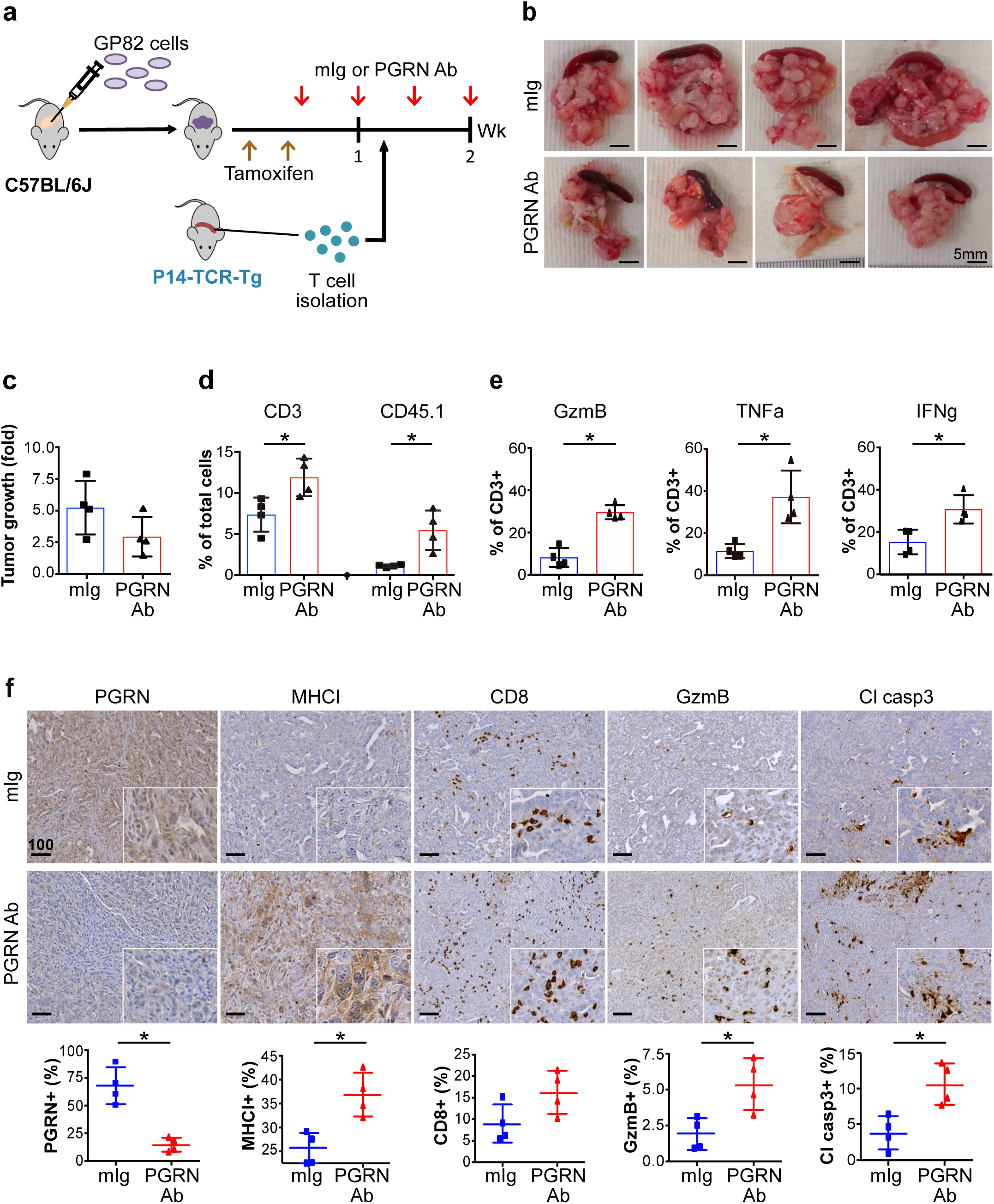
*In vivo* PGRN blockade promotes antigen-specific T cell cytotoxicity against tumors in orthotopic FKPC2GP model. **(a)** Timeline for treatment of anti-PGRN antibody (PGRN Ab) or mIg (50mg/kg) in orthotopic model of GP82 cells in C57BL/6J mice. **(b)** Tumors and spleens of mIg-treated (n=4) and PGRN Ab-treated (n=4) mice with orthotopic GP82 transplantation and intravenous injection of LCMV-gp33-reactive T cells freshly isolated from P14-TCR-Tg mice. **(c)** Tumor growth was assessed by ultrasound imaging and presented as fold change in tumor volume before and after PGRN Ab or mIg treatment started. **(d)** Tumors were digested into disaggregated cells and stained for T cell infiltration. Flow cytometric analysis showing the percentage of tumor infiltrating T (CD3+) cells and the exogenously injected LCMV-gp33-reactive (CD45.1+) T cells, in tumors treated with PGRN Ab (n=4) or mIg (n=4). **(e)** Flow cytometric analysis showing the expression of cytotoxic markers granzyme B (GzmB), TNFa and IFNg on CD3+ T cells in tumors with PGRN Ab (n=4) or mIg (n=4). **(f)** IHC staining of PGRN, MHCI marker H-2Db, CD8, GzmB and cleaved caspase 3 (Cl Casp3) in orthotopic GP82 tumors with PGRN Ab (n=4) or mIg (n=4). The lower panels show the percentages of positive cells in the whole tumors quantified by Definiens. Mean <u>±</u> SD is shown. *p<0.05. Scale bar unit: µm

LCMV-gp33 expression in the tumors was confirmed, using the GP82 xenografts that were not treated with tamoxifen as controls **(Figure S8a,b)**. Strikingly, tumor size was prominently reduced upon PGRN Ab treatment when compared to mIg **(Figure 7b)**. The progression of tumor volume was monitored by ultrasound imaging. Tumor increase in PGRN Ab treated tumor was slightly lower than that of mIg treatment, though not statistically significant due to limited subject numbers **(Figure 7c)**. Importantly however, total T cell (CD3+) and exogenous LCMV-gp33-reactive T cell (CD45.1+) infiltration were both increased upon PGRN Ab treatment **(Figure 7d)**. Furthermore, levels of GzmB, TNFa and IFNg were all significantly increased in PGRN Ab-treated tumors **(Figure 7e)**. IHC confirmed that in PGRN Ab-treated mice, PGRN levels were reduced while MHCI, CD8+ T cell infiltration, GzmB and cleaved caspase3 levels were all substantially increased compared to controls **(Figure 7f)**. Notably, in control mice without LCMV-gp33 expression and PGRN Ab, injection of LCMV-gp33-reactive T cells did not elicit any CD8 infiltration and activity **(Figure S8c)**. As a further control, treatment with both LCMV-gp33 expression and PGRN Ab treatment, but without LCMV-gp33-reactive T cell injection, showed no significant CD8 infiltration and anti-tumor cytotoxicity, and no prominent benefit in tumor growth suppression **(Figure S8d,e)**. Since functional macrophages are present, the results imply that the effect of PGRN Ab on macrophages as shown earlier does not contribute prominently to the anti-tumor cytotoxicity in this model system. Overall, we conclude that PGRN blockade promotes antigen-specific T cell cytotoxicity in endogenous PDAC.

## DISCUSSION

Understanding the mechanisms exploited by the tumors is critical to overcome immune evasion. PDAC has relatively few coding mutations, and thus few neoantigenic targets. In light of this, antigenic tumor peptides or dendritic cells loaded with shared peptides have been recently introduced to the clinic. However, despite the activation of specific anti-tumor T cell immunity, the observed tumor regressions are so far below expectations, and one of the possible reasons could be the absence or low level of tumor MHCI expression. Here, we unveiled an imperative role of PGRN in tumor cells as a key instructive regulator of immune evasion via MHCI regulation in PDAC cells in cell-autonomous manner.

We provided evidence that PGRN blockade restored MHCI expression by inhibiting autophagy. A role for autophagy in MHCI downregulation in pancreatic cancer was recently demonstrated by Yamamoto *et al* (31). In addition to autophagy, there are various mechanisms regulating MHCI expression in tumor cells, such as NF**-**kB stabilization as well as IFN, STAT3, and TGF-β signaling. Interestingly, the TGF-β gene signature was significantly enriched in *GRN*-high tumors of the Maurer *et al* dataset (19), implying TGF-β signaling as another potential mechanism underlying PGRN-induced MHCI downregulation in PDAC. Besides classical MHCI, PGRN is also known to decrease expression of the non-classical MHCI molecule MICA on tumor cells (16, 17), recently reported to impair co-stimulation of CD8 T cells (38). Investigation of the secretome from PGRN-expressing tumor cells could provide further insights into additional PGRN-mediated immunosuppressive constraints in PDAC, and potentially aid in stratification of tumors for appropriate treatment strategies.

PGRN blockade with Ab was used *in vivo* to demonstrate the functional effect of PGRN. Since PGRN is expressed in both tumor cells and macrophages, where different biological effects are exerted, our Ab approach is not able to distinguish the functions and significance of PGRN in the two compartments. However, from a translational perspective, neutralizing the secretory PGRN and subsequently reducing cellular PGRN to achieve systemic PGRN blockage by PGRN Ab treatment is of value for future therapeutic development. Indeed, in addition to MHCI regulation, we anticipate that PGRN might be involved in other immune evasion mechanisms. Analysis from the Maurer *et al* dataset supports an enrichment of mesenchymal phenotypes in high *GRN* epithelium, including upregulated EMT, TGF-β, glycolysis and down-regulation in oxidative phosphorylation. It is known that mesenchymal tumor cells possess high immunoevasive abilities (39–41). Further investigations are required to comprehensively delineate the signaling pathways underlying PGRN-mediated immunoevasion in tumor cells.

The role of macrophage-derived PGRN was well-characterized in liver metastasis of PDAC by Nielsen *et al*, who elegantly demonstrated that macrophage-derived PGRN activated resident hepatic stellate cells (HSCs) into myofibroblasts secreating periostin, resulting in a fibrotic microenvironment that promoted metastatic tumor growth and created physical barrier to CD8 infiltration(8, 9). However, in the CONKO-001 cohort, PGRN expression was not associated with α-SMA intensity **(Table S8)**. Besides, PGRN blockade *in vivo* also did not lead to reduction in fibroblast accumulation (α-SMA+ cells) in the tumors **(Figure S5D)**, suggesting the revived CD8 activity induced was not due to fibrosis reduction. One potential explanation might be the difference between pancreatic stellate cells (PSCs) and HSCs. Despite the similar morphological and functional features shared by the two, distinctive organ-specific differences were observed between their transcriptomes. When compared to HSCS, PSCs express higher levels of HIF1A, integrin α7, connective tissue growth factor, cytoskeletal elements, etc (42). Besides, the exact origin of PSCs is still controversial as these cells express both mesenchymal and neuroectodermal markers. Better understanding of PSCs is therefore needed in order to clarify the influence of PGRN on them in PDAC.

Our findings identify a potential key mechanism exploited by tumor cells for immune evasion in PDAC. Future studies on targeting PGRN in combination with other immunomodulatory therapeutic strategies are needed to explore its clinical significance in inducing more durable tumor rejection by targeting central immune escape routes in pancreatic cancer. Given the devastating failure of immunotherapy in PDAC and yet the exciting developments in chimeric antigen receptor T cell development, PGRN may be a pivotal target to enhance potent tumor antigen-mediated cytotoxicity.

## Supporting information

Supplementary Experimental Procedures

## ACKNOWLEDGMENTS

The authors would like to thank Jacks T and Tuveson DA for providing the KrasLSL-G12D mice; Nakhai H and Schmid RM for the PTF1A-Cre; Berns A for the lox p53 mice; Saur D for the Rosa-Cre-ERTKras^FSF-G12D^ mice; Kirsch DG for the frt p53 mice; Zinkernagel RM for the *Tg(TcrLCMV)^327Sdz^* mice.

## REFERENCES

1. Nguyen KB, Spranger S. Modulation of the immune microenvironment by tumor-intrinsic oncogenic signaling. The Journal of cell biology 2020;219(1) doi 10.1083/jcb.201908224.

2. Gonzalez H, Hagerling C, Werb Z. Roles of the immune system in cancer: from tumor initiation to metastatic progression. Genes & development 2018;32(19-20):1267–84 doi 10.1101/gad.314617.118.

3. Garrido F, Aptsiauri N, Doorduijn EM, Garcia Lora AM, van Hall T. The urgent need to recover MHC class I in cancers for effective immunotherapy. Current opinion in immunology 2016;39:44–51 doi 10.1016/j.coi.2015.12.007.

4. Beatty GL, Gladney WL. Immune escape mechanisms as a guide for cancer immunotherapy. Clinical cancer research : an official journal of the American Association for Cancer Research 2015;21(4):687–92 doi 10.1158/1078-0432.CCR-14-1860.

5. McGranahan N, Rosenthal R, Hiley CT, Rowan AJ, Watkins TBK, Wilson GA, et al. Allele-Specific HLA Loss and Immune Escape in Lung Cancer Evolution. Cell 2017;171(6):1259–71 e11 doi 10.1016/j.cell.2017.10.001.

6. Such L, Zhao F, Liu D, Thier B, Le-Trilling VTK, Sucker A, et al. Targeting the innate immunoreceptor RIG-I overcomes melanoma-intrinsic resistance to T cell immunotherapy. The Journal of clinical investigation 2020;130(8):4266–81 doi 10.1172/JCI131572.

7. Rodig SJ, Gusenleitner D, Jackson DG, Gjini E, Giobbie-Hurder A, Jin C, et al. MHC proteins confer differential sensitivity to CTLA-4 and PD-1 blockade in untreated metastatic melanoma. Sci Transl Med 2018;10(450) doi 10.1126/scitranslmed.aar3342.

8. Nielsen SR, Quaranta V, Linford A, Emeagi P, Rainer C, Santos A, et al. Macrophage-secreted granulin supports pancreatic cancer metastasis by inducing liver fibrosis. Nature cell biology 2016;18(5):549–60 doi 10.1038/ncb3340.

9. Quaranta V, Rainer C, Nielsen SR, Raymant ML, Ahmed MS, Engle DD, et al. Macrophage-Derived Granulin Drives Resistance to Immune Checkpoint Inhibition in Metastatic Pancreatic Cancer. Cancer research 2018;78(15):4253–69 doi 10.1158/0008-5472.CAN-17-3876.

10. Cenik B, Sephton CF, Kutluk Cenik B, Herz J, Yu G. Progranulin: a proteolytically processed protein at the crossroads of inflammation and neurodegeneration. The Journal of biological chemistry 2012;287(39):32298–306 doi 10.1074/jbc.R112.399170.

11. Cheung ST, Cheung PF, Cheng CK, Wong NC, Fan ST. Granulin-epithelin precursor and ATP-dependent binding cassette (ABC)B5 regulate liver cancer cell chemoresistance. Gastroenterology 2011;140(1):344–55 doi 10.1053/j.gastro.2010.07.049.

12. Cheung ST, Wong SY, Leung KL, Chen X, So S, Ng IO, et al. Granulin-epithelin precursor overexpression promotes growth and invasion of hepatocellular carcinoma. Clinical cancer research : an official journal of the American Association for Cancer Research 2004;10(22):7629–36 doi 10.1158/1078-0432.CCR-04-0960.

13. Yang D, Wang LL, Dong TT, Shen YH, Guo XS, Liu CY, et al. Progranulin promotes colorectal cancer proliferation and angiogenesis through TNFR2/Akt and ERK signaling pathways. American journal of cancer research 2015;5(10):3085–97.

14. Lu R, Serrero G. Mediation of estrogen mitogenic effect in human breast cancer MCF-7 cells by PC-cell-derived growth factor (PCDGF/granulin precursor). Proceedings of the National Academy of Sciences of the United States of America 2001;98(1):142–7 doi 10.1073/pnas.011525198.

15. Monami G, Gonzalez EM, Hellman M, Gomella LG, Baffa R, Iozzo RV, et al. Proepithelin promotes migration and invasion of 5637 bladder cancer cells through the activation of ERK1/2 and the formation of a paxillin/FAK/ERK complex. Cancer research 2006;66(14):7103–10 doi 10.1158/0008-5472.CAN-06-0633.

16. Cheung PF, Yip CW, Ng LW, Wong CK, Cheung TT, Lo CM, et al. Restoration of natural killer activity in hepatocellular carcinoma by treatment with antibody against granulin-epithelin precursor. Oncoimmunology 2015;4(7):e1016706 doi 10.1080/2162402X.2015.1016706.

17. Cheung PF, Yip CW, Wong NC, Fong DY, Ng LW, Wan AM, et al. Granulin-epithelin precursor renders hepatocellular carcinoma cells resistant to natural killer cytotoxicity. Cancer immunology research 2014;2(12):1209–19 doi 10.1158/2326-6066.CIR-14-0096.

18. Oettle H, Neuhaus P, Hochhaus A, Hartmann JT, Gellert K, Ridwelski K, et al. Adjuvant chemotherapy with gemcitabine and long-term outcomes among patients with resected pancreatic cancer: the CONKO-001 randomized trial. Jama 2013;310(14):1473–81 doi 10.1001/jama.2013.279201.

19. Maurer C, Holmstrom SR, He J, Laise P, Su T, Ahmed A, et al. Experimental microdissection enables functional harmonisation of pancreatic cancer subtypes. Gut 2019;68(6):1034–43 doi 10.1136/gutjnl-2018-317706.

20. Cima I, Kong SL, Sengupta D, Tan IB, Phyo WM, Lee D, et al. Tumor-derived circulating endothelial cell clusters in colorectal cancer. Sci Transl Med 2016;8(345):345ra89 doi 10.1126/scitranslmed.aad7369.

21. Schug J, Schuller WP, Kappen C, Salbaum JM, Bucan M, Stoeckert CJ, Jr. Promoter features related to tissue specificity as measured by Shannon entropy. Genome Biol 2005;6(4):R33 doi 10.1186/gb-2005-6-4-r33.

22. Tan WJ, Cima I, Choudhury Y, Wei X, Lim JC, Thike AA, et al. A five-gene reverse transcription-PCR assay for pre-operative classification of breast fibroepithelial lesions. Breast Cancer Res 2016;18(1):31 doi 10.1186/s13058-016-0692-6.

23. 23. Baddeley AR, E.; Turner, R. Spatial Point Patterns: Methodology and Applications with R.: CRC Press; 2015.

24. Mazur PK, Herner A, Mello SS, Wirth M, Hausmann S, Sanchez-Rivera FJ, et al. Combined inhibition of BET family proteins and histone deacetylases as a potential epigenetics-based therapy for pancreatic ductal adenocarcinoma. Nature medicine 2015;21(10):1163–71 doi 10.1038/nm.3952.

25. Page N, Klimek B, De Roo M, Steinbach K, Soldati H, Lemeille S, et al. Expression of the DNA-Binding Factor TOX Promotes the Encephalitogenic Potential of Microbe-Induced Autoreactive CD8(+) T Cells. Immunity 2018;48(5):937–50 e8 doi 10.1016/j.immuni.2018.04.005.

26. Cheung PF, Cheng CK, Wong NC, Ho JC, Yip CW, Lui VC, et al. Granulin-epithelin precursor is an oncofetal protein defining hepatic cancer stem cells. PloS one 2011;6(12):e28246 doi 10.1371/journal.pone.0028246.

27. Ho JC, Ip YC, Cheung ST, Lee YT, Chan KF, Wong SY, et al. Granulin-epithelin precursor as a therapeutic target for hepatocellular carcinoma. Hepatology 2008;47(5):1524–32 doi 10.1002/hep.22191.

28. Groeneveldt C, van Hall T, van der Burg SH, Ten Dijke P, van Montfoort N. Immunotherapeutic Potential of TGF-beta Inhibition and Oncolytic Viruses. Trends in immunology 2020;41(5):406–20 doi 10.1016/j.it.2020.03.003.

29. Lohneis P, Sinn M, Bischoff S, Juhling A, Pelzer U, Wislocka L, et al. Cytotoxic tumour-infiltrating T lymphocytes influence outcome in resected pancreatic ductal adenocarcinoma. European journal of cancer 2017;83:290–301 doi 10.1016/j.ejca.2017.06.016.

30. Striefler JK, Sinn M, Pelzer U, Juhling A, Wislocka L, Bahra M, et al. P53 overexpression and Ki67-index are associated with outcome in ductal pancreatic adenocarcinoma with adjuvant gemcitabine treatment. Pathology, research and practice 2016;212(8):726–34 doi 10.1016/j.prp.2016.06.001.

31. Yamamoto K, Venida A, Yano J, Biancur DE, Kakiuchi M, Gupta S, et al. Autophagy promotes immune evasion of pancreatic cancer by degrading MHC-I. Nature 2020;581(7806):100–5 doi 10.1038/s41586-020-2229-5.

32. Tanaka Y, Suzuki G, Matsuwaki T, Hosokawa M, Serrano G, Beach TG, et al. Progranulin regulates lysosomal function and biogenesis through acidification of lysosomes. Hum Mol Genet 2017;26(5):969–88 doi 10.1093/hmg/ddx011.

33. Elia LP, Mason AR, Alijagic A, Finkbeiner S. Genetic Regulation of Neuronal Progranulin Reveals a Critical Role for the Autophagy-Lysosome Pathway. J Neurosci 2019;39(17):3332–44 doi 10.1523/JNEUROSCI.3498-17.2019.

34. Chang MC, Srinivasan K, Friedman BA, Suto E, Modrusan Z, Lee WP, et al. Progranulin deficiency causes impairment of autophagy and TDP-43 accumulation. The Journal of experimental medicine 2017;214(9):2611–28 doi 10.1084/jem.20160999.

35. Beel S, Moisse M, Damme M, De Muynck L, Robberecht W, Van Den Bosch L, et al. Progranulin functions as a cathepsin D chaperone to stimulate axonal outgrowth in vivo. Hum Mol Genet 2017;26(15):2850–63 doi 10.1093/hmg/ddx162.

36. Wong NC, Cheung PF, Yip CW, Chan KF, Ng IO, Fan ST, et al. Antibody against granulin-epithelin precursor sensitizes hepatocellular carcinoma to chemotherapeutic agents. Molecular cancer therapeutics 2014;13(12):3001–12 doi 10.1158/1535-7163.MCT-14-0012.

37. Cheung PF, Neff F, Neander C, Bazarna A, Savvatakis K, Liffers ST, et al. Notch-induced myeloid reprogramming in spontaneous pancreatic ductal adenocarcinoma by dual genetic targeting. Cancer research 2018 doi 10.1158/0008-5472.CAN-18-0052.

38. Groh V, Wu J, Yee C, Spies T. Tumour-derived soluble MIC ligands impair expression of NKG2D and T-cell activation. Nature 2002;419(6908):734–8 doi 10.1038/nature01112.

39. Lou Y, Diao L, Cuentas ER, Denning WL, Chen L, Fan YH, et al. Epithelial-Mesenchymal Transition Is Associated with a Distinct Tumor Microenvironment Including Elevation of Inflammatory Signals and Multiple Immune Checkpoints in Lung Adenocarcinoma. Clinical cancer research : an official journal of the American Association for Cancer Research 2016;22(14):3630–42 doi 10.1158/1078-0432.CCR-15-1434.

40. Alsuliman A, Colak D, Al-Harazi O, Fitwi H, Tulbah A, Al-Tweigeri T, et al. Bidirectional crosstalk between PD-L1 expression and epithelial to mesenchymal transition: significance in claudin-low breast cancer cells. Mol Cancer 2015;14:149 doi 10.1186/s12943-015-0421-2.

41. Tamborero D, Rubio-Perez C, Muinos F, Sabarinathan R, Piulats JM, Muntasell A, et al. A Pan-cancer Landscape of Interactions between Solid Tumors and Infiltrating Immune Cell Populations. Clinical cancer research : an official journal of the American Association for Cancer Research 2018;24(15):3717–28 doi 10.1158/1078-0432.CCR-17-3509.

42. Buchholz M, Kestler HA, Holzmann K, Ellenrieder V, Schneiderhan W, Siech M, et al. Transcriptome analysis of human hepatic and pancreatic stellate cells: organ-specific variations of a common transcriptional phenotype. J Mol Med (Berl) 2005;83(10):795–805 doi 10.1007/s00109-005-0680-2.

