## Supplementary Figures S1-8 for "Progranulin promotes immune evasion of pancreatic adenocarcinoma through regulation of MHCI expression"

### Figure S1

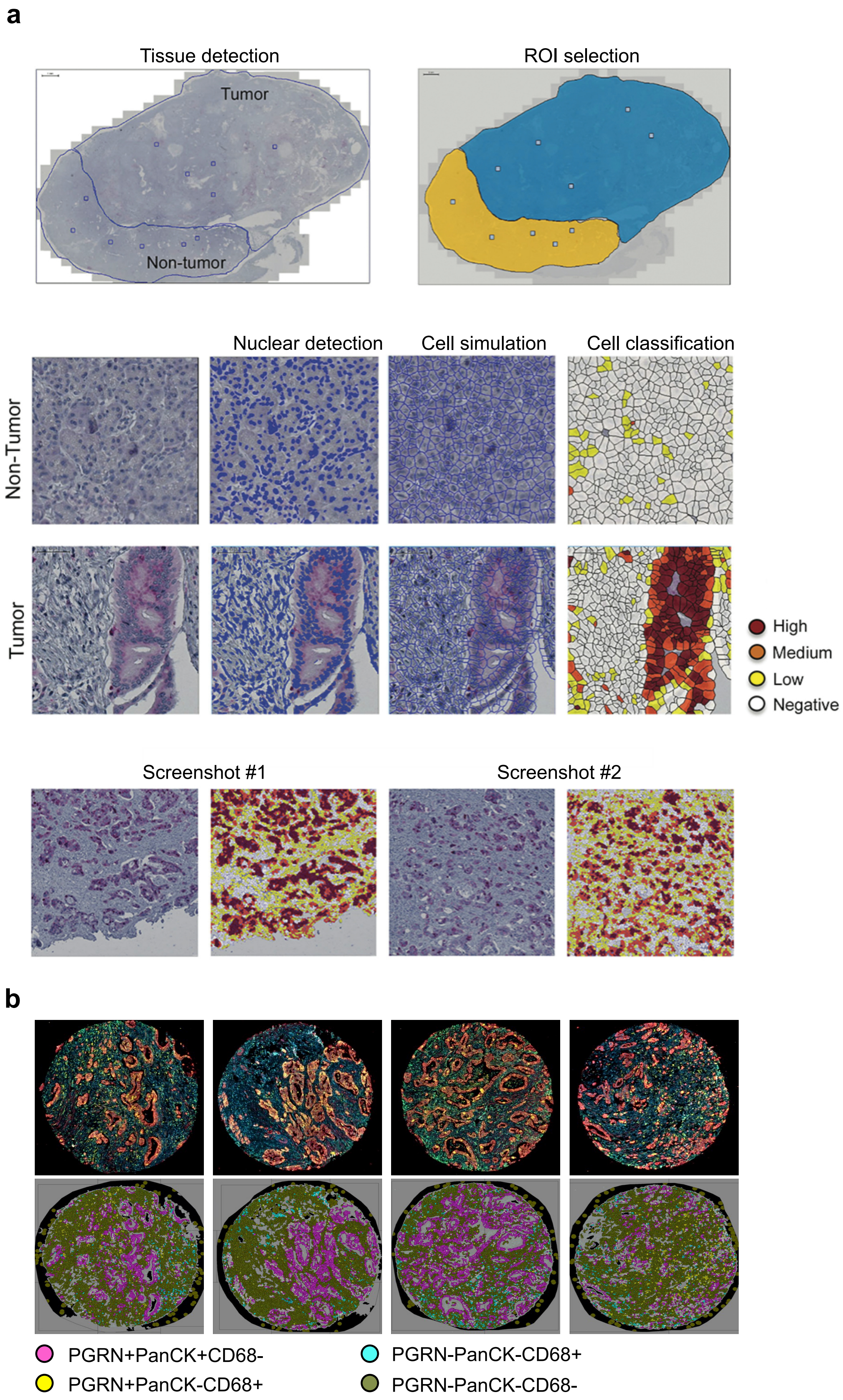
a
Tissue detection
Tumor
Non-tumor
ROI selection
Nuclear detection
Cell simulation
Cell classification
Non-Tumor
Tumor
High
Medium
Low
Negative
Screenshot #1
Screenshot #2
b
PGRN+PanCK+CD68-
PGRN-PanCK-CD68+
PGRN+PanCK-CD68+
PGRN-PanCK-CD68-

**Figure S1**

**(a)** Quantification of IHC staining. Tumor and non-tumor pancreas tissue were defined. Regions of interest (ROIs) were selected for defining nucleus and cells, for accurate assessment of cell number of each slide. Cell classification was performed to categorized different signal intensity. Screenshot #1 and #2 as examples to illustrate the accuracy of the software. **(b)** Quantification of mIF staining in TMA. Examples showing quantification of cells co-expressing PGRN and/or PanCK and/or CD68.

### Figure S2

#### Essen cohort

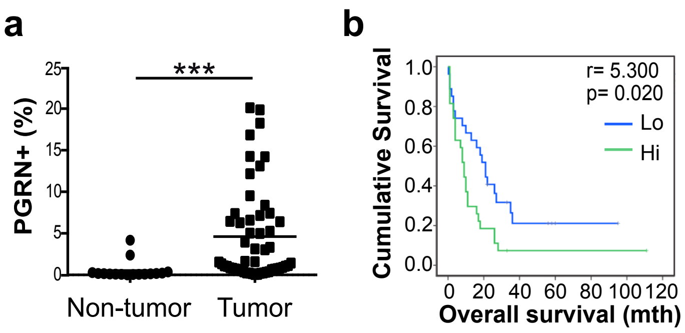


#### Nijmegen cohort

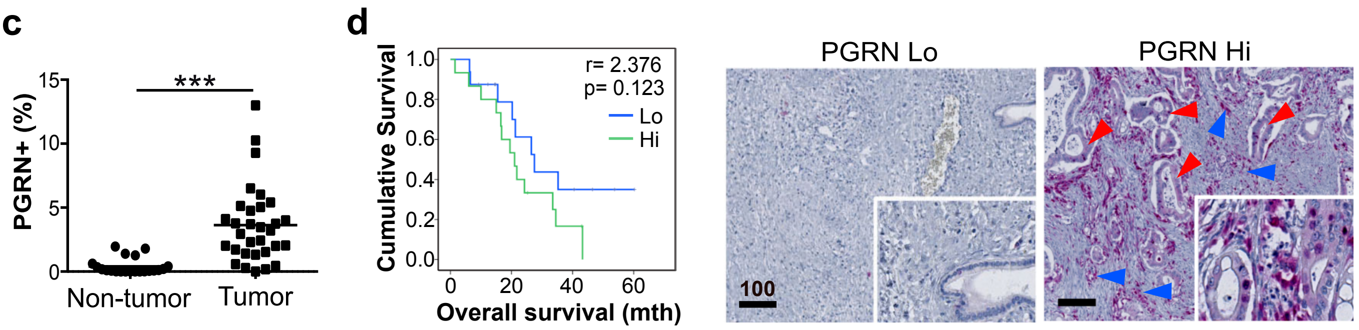


#### Maurer *et al* (GSE93326)

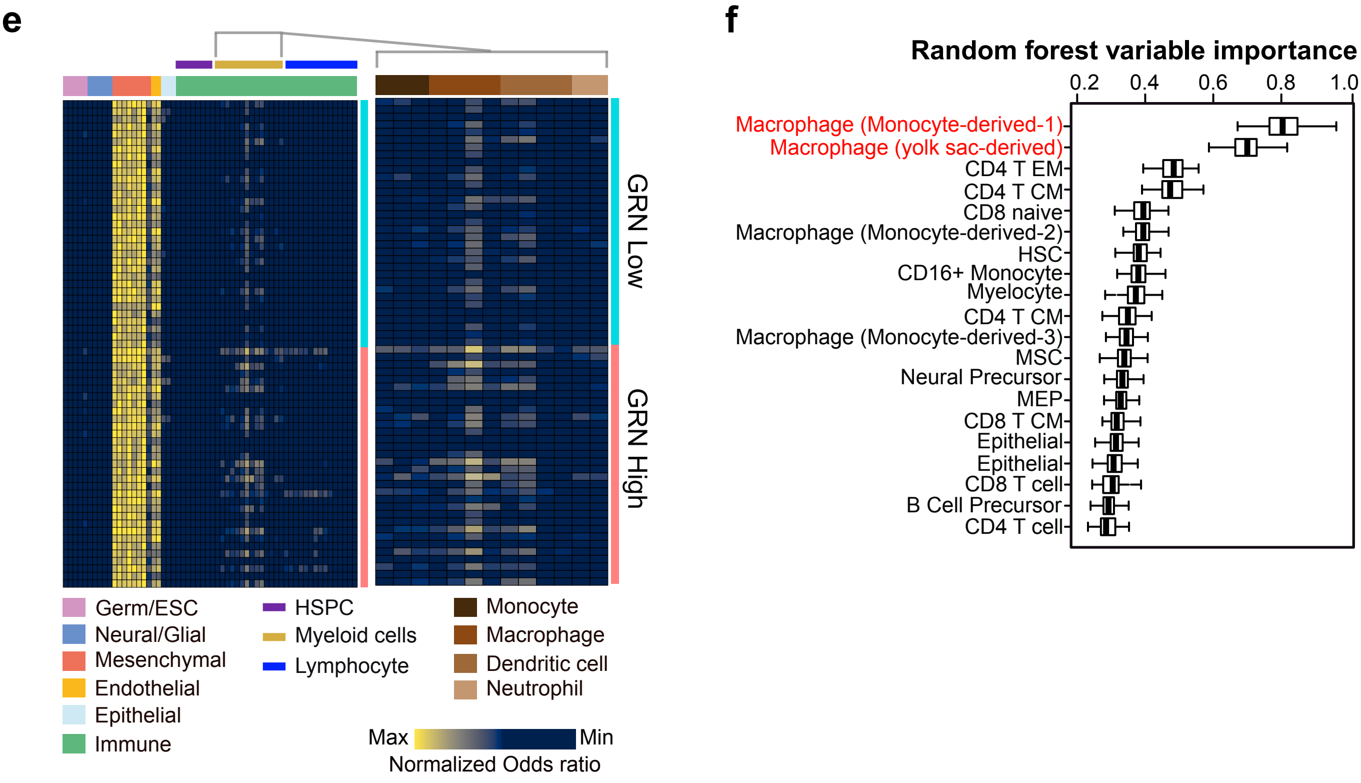

**Figure S2**

**(a,b**) Essen cohort. **(a)** PGRN+ cells in tumor (n=53) and adjacent non-tumor (n=16) tissues of human PDAC was assessed by IHC staining and quantified by Definiens. **(b)** Kaplan-Meier overall survival plot according to PGRN expression level. Patients were segregated into low (n=26) and high (n=27) expression groups with median of number of PGRN+ cells as cutoff. **(c,d)** Nijmegen cohort (n=31). **(c)** PGRN expression levels in tumor (n=31) and adjacent non-tumor (n=21) tissues of human PDAC were assessed by IHC staining and quantified by Definiens. **(d)** Kaplan-Meier overall survival plots according to PGRN expression level. Patients (n= 31) were segregated into low (n=16) and high (n=15) expression groups with median of positive cells as cutoff. Right panel shows representative IHC staining of high- and low- PGRN expressing patient specimens. ***Right panel:*** PGRN expression in both tumor and stromal compartments of PDAC. Red arrowheads indicate PGRN+ tumor cells; blue arrowheads indicate PGRN+ stromal cells. **(e,f)** Maurer *et al* dataset (GSE93326, n=65). **(e)** Cell type deconvolution of 43 different cell types of transcriptomes derived from *GRN*-high (*n* =32) and *GRN*-low (*n* =32) stroma samples, indicating enrichment of myeloid cells in *GRN*-high stroma. Data represents normalized odds ratios comparing number of enriched cell type-specific genes with random enrichment for each sample (rows) and cell type (columns). HSPC, Hematopoietic stem and progenitor cells. **(f)** Box plots represent top 20 variable importance computed by 100-fold random forest for each cell type identified in **(e)**, and ordered by descending importance in predicting *GRN* high and low stroma samples. Macrophages represents the cell type showing highest degree of importance in predicting *GRN* high stroma. *** p<0.001, Scale bar unit: µm

Figure S3

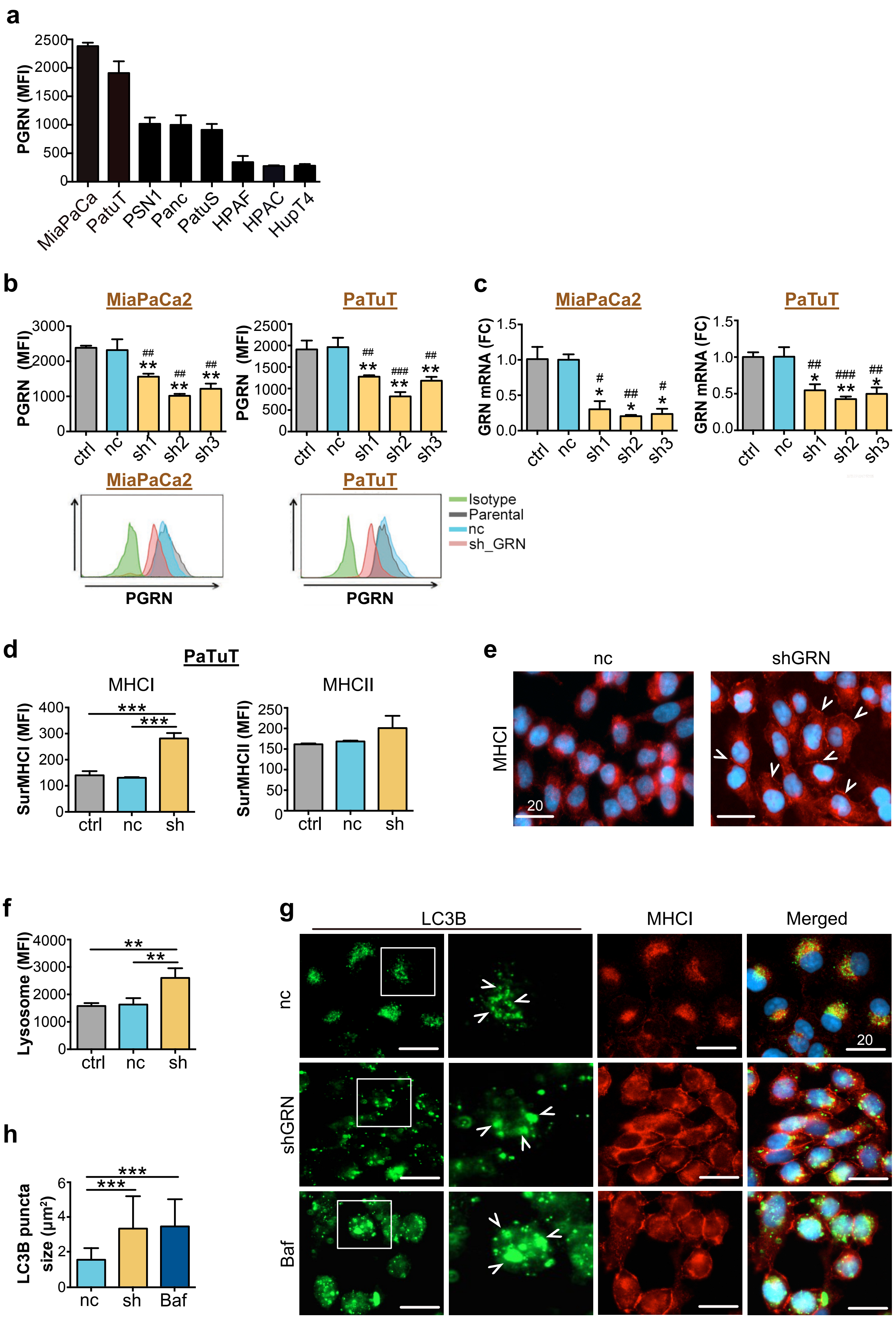
a
PGRN (MFI)
2500
2000
1500
1000
500
0
MiaPaCa
PatuT
PSN1
Panc
PatuS
HPAF
HPAC
HupT4
b
MiaPaCa2
PaTuT
PGRN (MFI)
3000
2000
1000
0
2500
1500
500
ctrl
nc
sh1
sh2
sh3
##
###
**
c
GRN mRNA (FC)
1.5
1.0
0.5
0.0
#
*
Isotype
Parental
nc
sh_GRN
PGRN
d
PaTuT
MHCI
MHCII
SurMHCI (MFI)
SurMHCII (MFI)
400
300
200
100
250
150
50
***
ctrl
nc
sh
e
nc
shGRN
MHCI
20
f
Lysosome (MFI)
4000
3000
2000
1000
**
g
LC3B
MHCI
Merged
nc
shGRN
Baf
20
h
LC3B puncta
size (μm²)
6
4
2
0
***
nc
sh
Baf

**Figure S3**

**(a)** Expression level of PGRN in human commercial PDAC cell lines was assessed by flow cytometric analysis. **(b,c)** *GRN* suppression by shRNA plasmid transfection in PDAC cell lines MiaPaCa2 and PaTuT. PGRN protein (PGRN) and gene (*GRN*) levels measured by **(b)** flow cytometry and **(c)** qRT-PCR, respectively. FC: fold change. ctrl: parental PDAC cells; nc: shRNA scrambled control; sh1-3; *GRN* shRNA1-3. *GRN* shRNA2 (sh2) consistently demonstrated the highest efficiency of suppression and was therefore selected for subsequent functional evaluation as "shGRN". **(d)** Surface MHCI (HLA-A/B/C) and MHCII (HLA-DR) expression on human PDAC cell line PatuT upon *GRN* suppression is assessed by flow cytometry. **(e)** IF staining of MHCI marker HLA-A/B/C in PatuT upon *GRN* suppression. White arrowheads indicate the membraneous staining of MHCI. **(f)** Lysosome content in PatuT upon GRN suppression is assessed by staining with fluorescent LysoGreen Indicator and measured by flow cytometry. **(g, h)** IF staining of MHCI (red) and LC3B (green) in PatuT cells upon *GRN* suppression or treated with autophagy inhibitor Bafinomycin (100nM, 24h). **(h)** Size of LC3B puncta of 30 cells per treatment was measured by ZEN software. ctrl: parental PDAC cells; nc: shRNA scrambled control; sh/shGRN: *GRN* shRNA; Baf: Bafinomycin. Mean ± SD is shown; MFI: mean fluorescence intensity. *$p < 0.05$; ** $p<0.01$; *** $p<0.001$; # $p<0.05$; ##$p<0.01$; ###$p<0.001$. Scale bar unit: µm

Figure S4

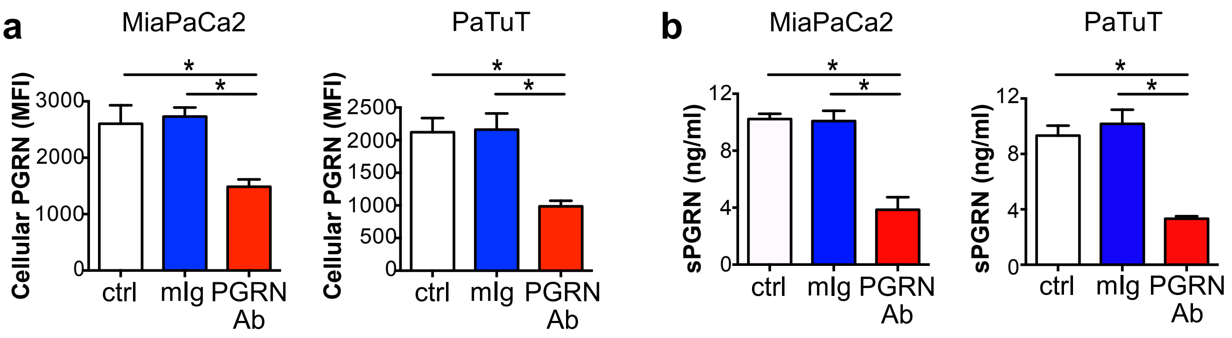

a
MiaPaCa2
PaTuT
Cellular PGRN (MFI)
3000
2000
1000
0
2500
2000
1500
1000
500
0
*
*
ctrl
mIg
PGRN Ab
b
MiaPaCa2
PaTuT
sPGRN (ng/ml)
12
8
4
0

**Figure S4**
MiaPaCa2 and PaTuT cells were serum-starved (1% FBS-supplemented medium) overnight, and then treated with PGRN Ab or mIg (100ug/ml) for 24h. Medium was then changed and cells were cultured for 2 more days. **(a)** Cellular PGRN level was assessed by flow cytometry, while **(b)** soluble PGRN (sPGRN) level was measured by ELISA. MFI: Mean Fluorescent Intensity. * $p<0.05$.

Figure S5

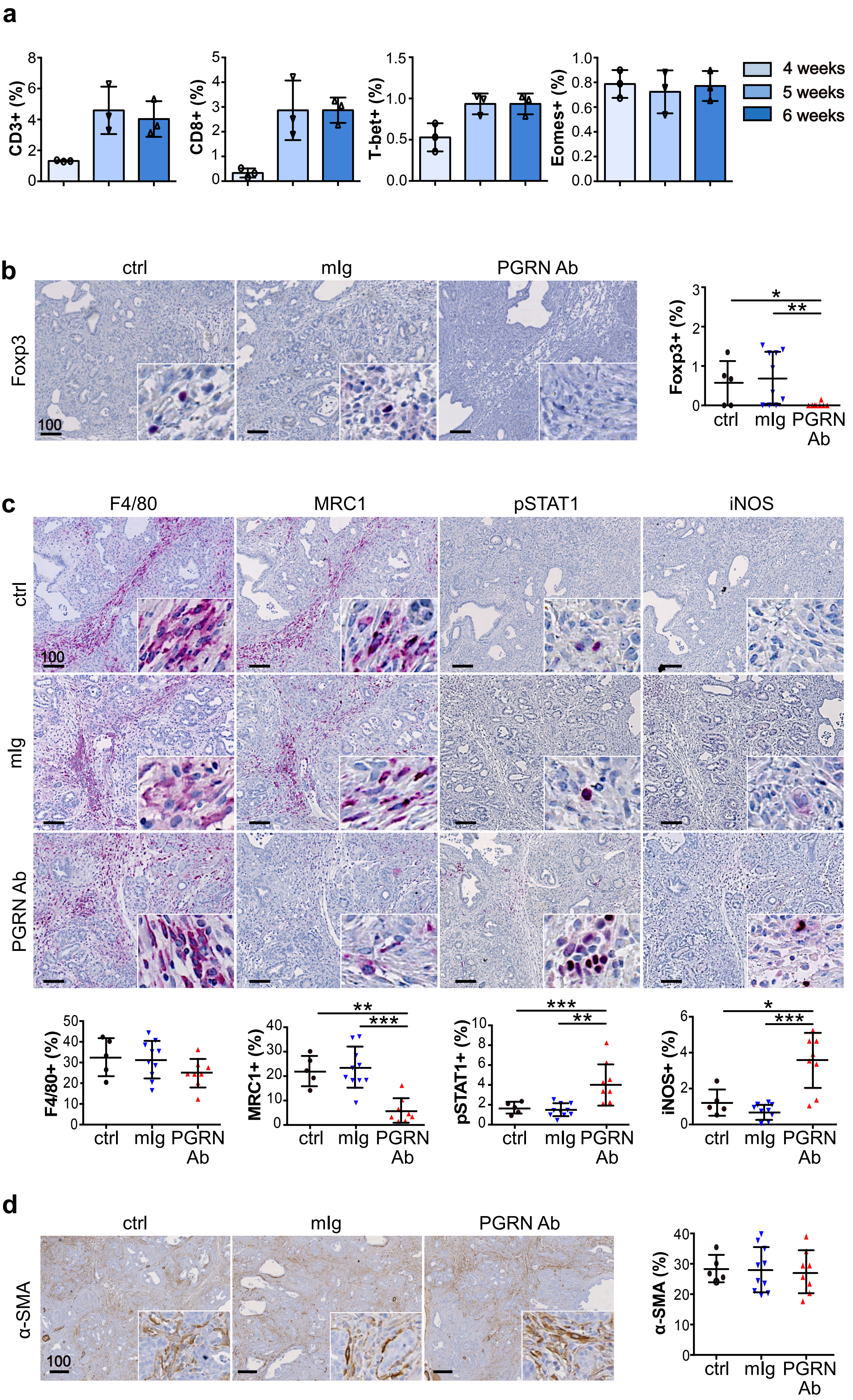
a
CD3+ (%)
CD8+ (%)
T-bet+ (%)
Eomes+ (%)
4 weeks
5 weeks
6 weeks
b
ctrl
mIg
PGRN Ab
Foxp3
100
Foxp3+ (%)
c
F4/80
MRC1
pSTAT1
iNOS
ctrl
mIg
PGRN Ab
F4/80+ (%)
MRC1+ (%)
pSTAT1+ (%)
iNOS+ (%)
d
α-SMA
α-SMA (%)

**Figure S5**
**(a**) Quantification of T cell markers CD3, CD8, T-bet and Eomes in *CKP* tumors during early PDAC development (n=3 per time point). **(b)** IHC of Foxp3 in *CKP* tumors treated with or without PGRN Ab or mIg. ctrl: n=5; mIg: n=10; PGRN Ab: n=8.The right panel shows the percentage of positive cells in the whole tumorous tissues. **(c)** IHC staining of pan-macrophage marker F4/80, M2 marker MRC1, M1 markers pSTAT1 and iNOS in *CKP* tumors treated with or without PGRN Ab or mIg (50mg/kg). ctrl: n=5; mIg: n=10; PGRN Ab: n=8. The lower panels show the percentage of positive cells in the whole tumors. **(d)** IHC staining of myofibroblast marker α-sma in *CKP* tumors treated with or without PGRN Ab (50mg/kg). ctrl: n=5; PGRN Ab: n=8. The right panel shows the percentage of α-SMA+cells in the whole tumors. *p< 0.05; ** p<0.01; *** p<0.001. Scale bar unit: μm

Figure S6

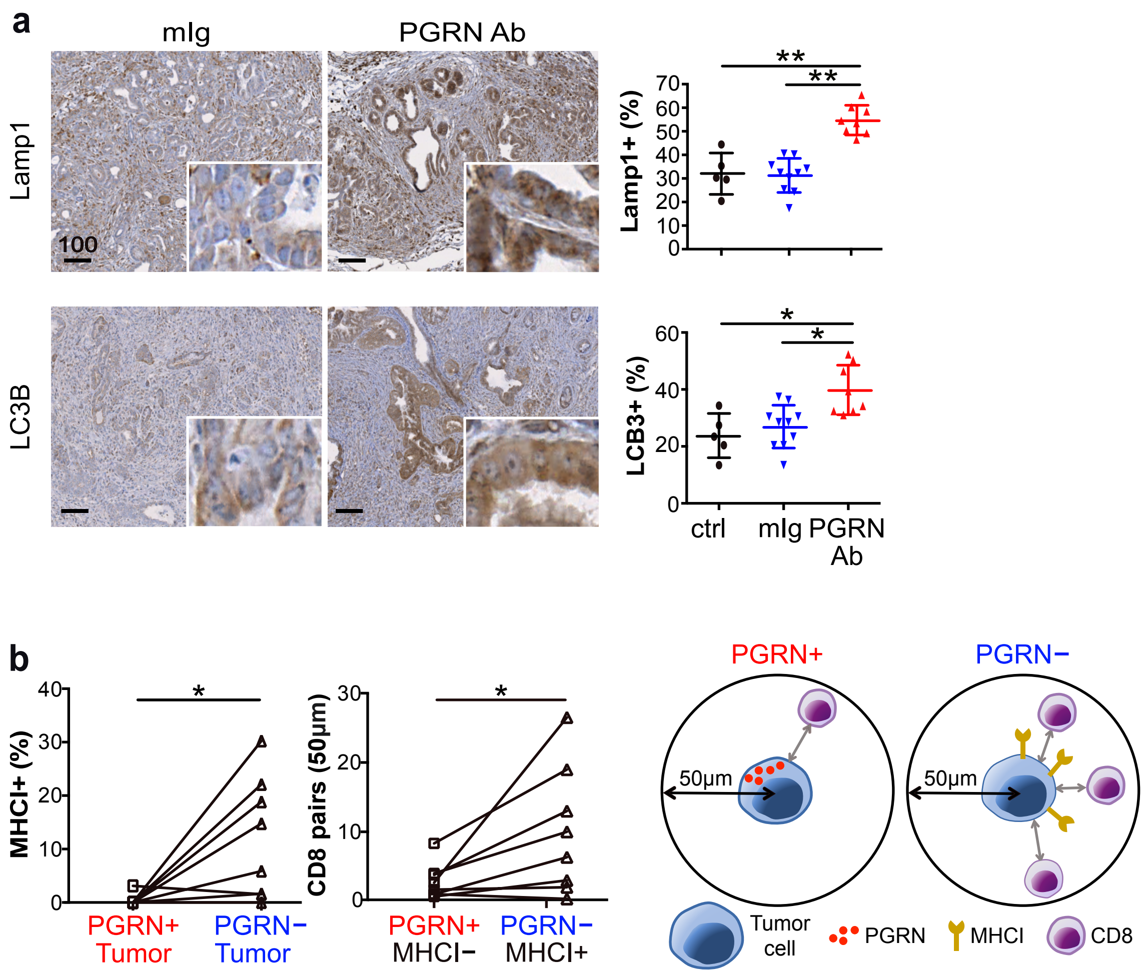
a
mIg
PGRN Ab
Lamp1
LC3B
100
Lamp1+ (%)
LCB3+ (%)
ctrl
mIg
PGRN Ab
b
MHCI+ (%)
PGRN+ Tumor
PGRN− Tumor
CD8 pairs (50µm)
PGRN+ MHCI−
PGRN− MHCI+
PGRN+
PGRN−
50µm
Tumor cell
PGRN
MHCI
CD8

**Figure S6**

**(a)** IHC staining of lysosome marker Lamp1 and autophagosome marker LC3B in *CKP* tumors treated with mIg (n=10) or PGRN Ab (n=8). The right panels show the percentage of Lamp1+ and LC3B+ cells in the whole tumorous tissues of *CKP* mice treated with or without PGRN Ab and mIg. **(b)** Automated computational analysis showing the percentage of MHCI+ cells in PGRN+/PanCK+ and PGRN-/PanCK+ populations (left panel), and the number of CD8+ cells in proximity (<50μm radical distance) of PGRN+/MHCI-/ PanCK+ or PGRN-/MHCI+/PanCK+ tumor cells in anti-PGRN Ab-treated *CKP* tumors (n=8) (right panel). Scheme depicts the differential MHCI expression and interaction of tumor and CD8 cells depending on tumor-expressed PGRN. *$p<0.05$; **$p<0.01$. Scale bar unit: μm

Figure S7

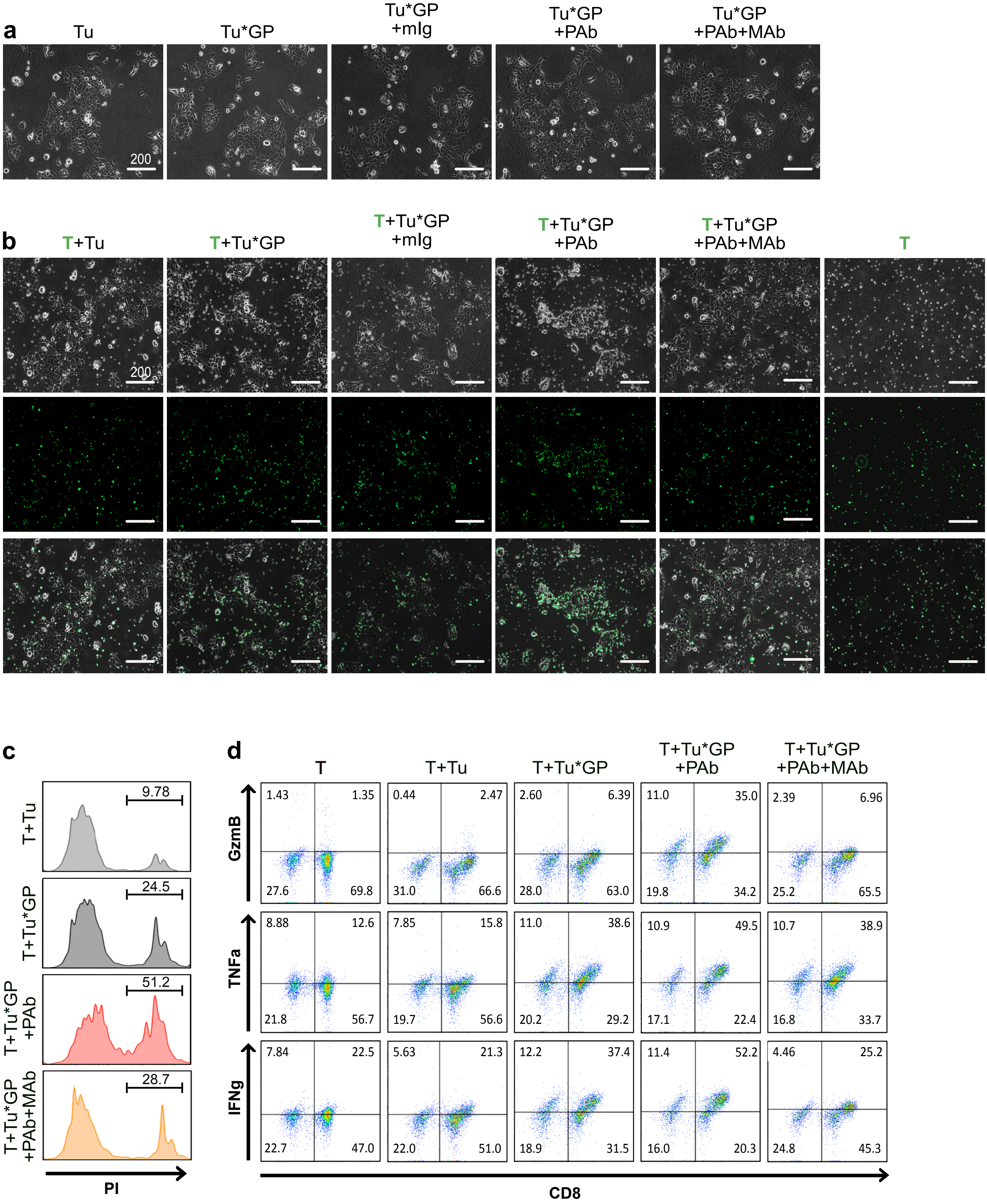
a
Tu
Tu*GP
Tu*GP +mIg
Tu*GP +PAb
Tu*GP +PAb+MAb
200
b
T+Tu
T+Tu*GP
T+Tu*GP +mIg
T+Tu*GP +PAb
T+Tu*GP +PAb+MAb
T
200
c
T+Tu
9.78
T+Tu*GP
24.5
T+Tu*GP +PAb
51.2
T+Tu*GP +PAb+MAb
28.7
PI
d
T
T+Tu
T+Tu*GP
T+Tu*GP +PAb
T+Tu*GP +PAb+MAb
GzmB
1.43 1.35 27.6 69.8
0.44 2.47 31.0 66.6
2.60 6.39 28.0 63.0
11.0 35.0 19.8 34.2
2.39 6.96 25.2 65.5
TNFa
8.88 12.6 21.8 56.7
7.85 15.8 19.7 56.6
11.0 38.6 20.2 29.2
10.9 49.5 17.1 22.4
10.7 38.9 16.8 33.7
IFNg
7.84 22.5 22.7 47.0
5.63 21.3 22.0 51.0
12.2 37.4 18.9 31.5
11.4 52.2 16.0 20.3
4.46 25.2 24.8 45.3
CD8

**Figure S7**

**(a)** Phase-contrast images of GP82 cells cultured with different treatments for 2 days. **(b)** Phase-contrast, fluorescence, and overlay images of GP82 cells and LCMV-gp33-reactive T cells (CFSE-labeled, green) after 2 days of co-culture. When anti-MHCI (H-2Db) neutralizing antibody (MAb) was included in the treatment, MAb was added 1h after PGRN Ab treatment. T cells were then added 1h after MAb treatment. Tu: Tumor cells (GP82 cells); GP: LCMV-gp33-induced; PAb: PGRN antibody (100ug/ml); MAb: MHCI (H-2Db) neutralizing antibody (100ug/ml). **(c)** Representative histograms showing cytotoxicity level of GP82 cells with or without LCMV-gp33 expression, anti-PGRN antibody, and anti-MHCI neutralizing antibody, upon co-culture with LCMV-gp33-reactive T cells. Percentages of PI+ cells are indicated. **(d)** Representative scatter plots showing the percentage of cells that are positive for both CD8 and cytotoxic markers granzyme B (GzmB), TNF-a, and IFN-g.

### Figure S8

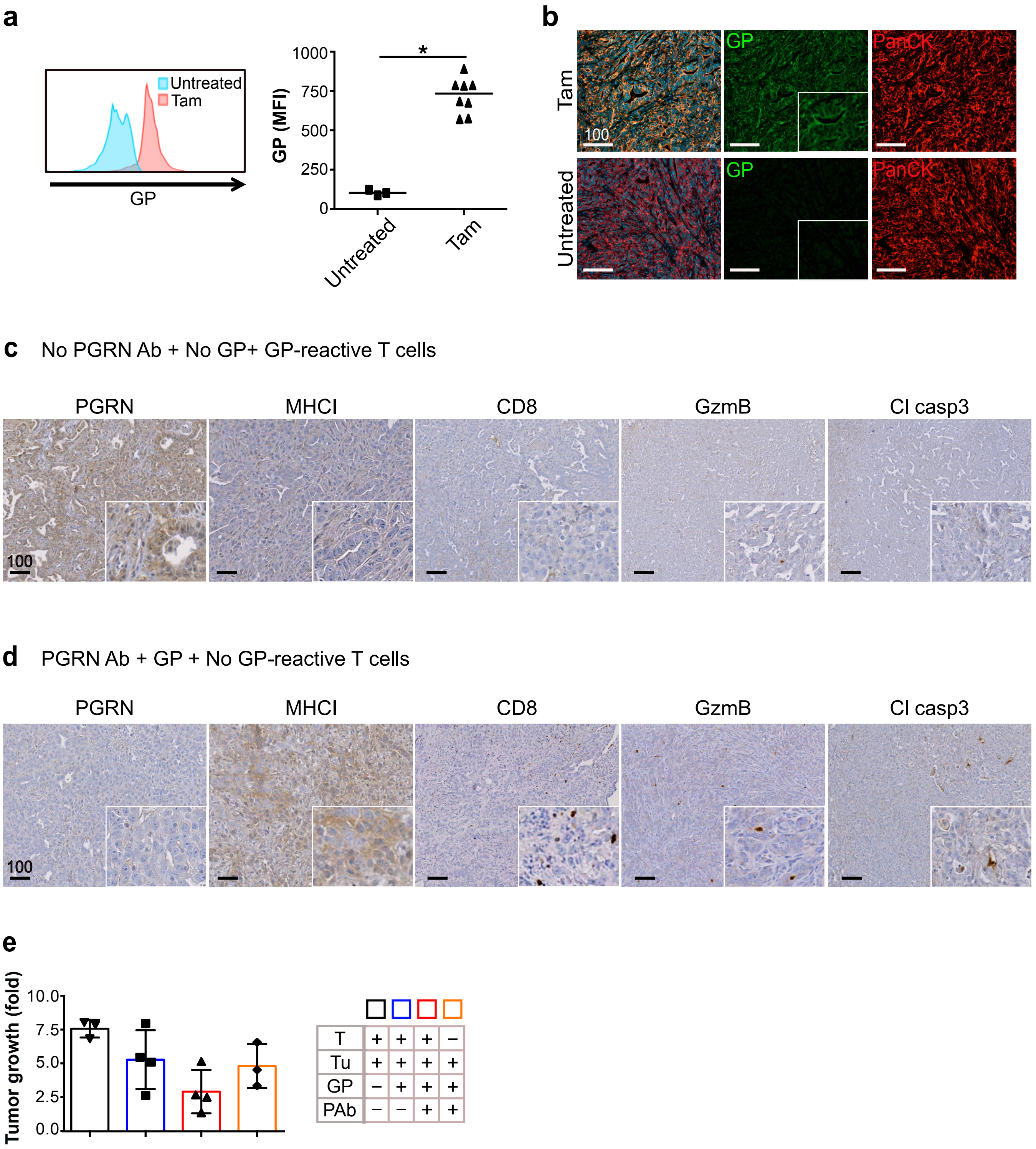
a
Untreated
Tam
GP
1000
750
500
250
0
GP (MFI)
*
Untreated
Tam
b
Tam
Untreated
GP
PanCK
100
c
No PGRN Ab + No GP+ GP-reactive T cells
PGRN
MHCI
CD8
GzmB
Cl casp3
100
d
PGRN Ab + GP + No GP-reactive T cells
PGRN
MHCI
CD8
GzmB
Cl casp3
100
e
Tumor growth (fold)
10.0
7.5
5.0
2.5
0.0
T + + + –
Tu + + + +
GP – + + +
PAb – – + +

**Figure S8**

**(a)** Tumors dissected from GP82 orthotopic models that are treated with or without tamoxifen (Tam, 75mg/kg) were digested into disaggregated cells. Cells were stained for LCMV-gp33 (GP) expression and analyzed by flow cytometry. **(b)** IF staining showing LCMV-gp33 (GP) expression in PanCK+ tumor cells in GP82 orthotopic models that were treated with or without tamoxifen (Tam, 75mg/kg). **(c, d)** IHC staining of PGRN, MHCI, CD8, GranzymB (GzmB), and cleaved casp3 (Cl casp3) in tumor **(c)** without tamoxifen and PGRN Ab treatment, but with LCMV-gp33-reactive T cell injection; and **(d)** with tamoxifen and PGRN Ab treatment, but without LCMV-gp33-reactive T cell injection. **(e)** Tumor growth was assessed by ultrasound imaging and presented as fold change in tumor volume before and after PGRN Ab or mIg treatment started. MFI: Mean Fluorescent Intensity.

*$p<0.05$. Scale bar unit: μm
