## Supplementary Table S2-6, S8 for "Progranulin promotes immune evasion of pancreatic adenocarcinoma through regulation of MHCI expression"

### Supplementary Tables: S2-6, 8

#### Table S2. IHC antibodies

| Antigen | Clone | Manufacturer | Dilution | Reactivity |
|---|---|---|---|---|
| PGRN | A23 | Ref [1] | 1:200 | Human, Mouse |
| PanCK | PCK-26 | Abcam | 1:100 | Human, Mouse |
| Ki67 | Polyclonal | Abcam | 1:200 | Mouse |
| F4/80 | BM8 | BMA biomedicals | 1:200 | Mouse |
| MRC1 | Polyclonal | Abcam | 1:200 | Mouse |
| INOS | Polyclonal | Abcam | 1:200 | Mouse |
| Phospho-STAT1 | M135 | Abcam | 1:100 | Mouse |
| Foxp3 | FJK-16s | ThermoFisher | 1:50 | Mouse |
| Cl. caspase 3 | 5A1E | Cell Signaling | 1:100 | Mouse |
| MHC I (H-2Db) | AF6-88.5.5.3 | ThermoFisher | 1:100 | Mouse |
| MHC II | M5/114.15.2 | ThermoFisher | 1:100 | Mouse |
| MHCI (HLA-A) | C-6 | Santa Cruz | 1:100 | Human |
| CD3 | Polyclonal | Abcam | 1:100 | Mouse |
| CD4 | RM4-5 | BD | 1:50 | |
| CD8 | SP16 | Abcam | 1:100 | Human |
| CD8 | EPR20305 | Abcam | 1:100 | Mouse |
| T-bet | 4B10 | eBioscience | 1:50 | Mouse |
| Eomes | Dan11mag | eBioscience | 1:50 | Mouse |
| Granzyme B | Polyclonal | Abcam | 1:500 (hu), 1:100 (ms) | Human, Mouse |
| $\alpha$-sma | Polyclonal | Abcam | 1:100 | Mouse |
| Lamp1 | Polyclonal | Abcam | 1:200 | Mouse |
| LC3B | Polyclonal | Abcam | 1:100 | Mouse |
| CD68 | KP1 | Abcam | 1:400 | Human |

**Table S3. Sequential multiplexed immunofluorescence staining protocol**

| Antigen | Primary antibody | | | TSA fluorophore | |
|---|---|---|---|---|---|
| | **Clone** | **Manufacturer** | **Dilution** | **Fluorophore** | **Dilution** |
| **Fig. 1** | | | | | |
| CD68 | KP1 | Abcam | 1:400 | Opal520 | 1:200 |
| PGRN | A23 | Ref [1] | 1:200 | Opal570 | 1:200 |
| PanCK | PCK-26 | Abcam | 1:100 | Opal780 | 1:100 |
| **Fig. 2** | | | | | |
| PGRN | A23 | Ref [1] | 1:200 | Opal570 | 1:200 |
| CD8 | SP16 | Abcam | 1:100 | Opal480 | 1:200 |
| GzmB | Polyclonal | Abcam | 1:500 | Opal690 | 1:200 |
| MHCI (HLA-A) | C-6 | Santa Cruz | 1:100 | Opal520 | 1:200 |
| PanCK | PCK-26 | Abcam | 1:100 | Opal780 | 1:100 |
| **Fig. 5** | | | | | |
| PGRN | A23 | Ref [1] | 1:200 | Opal570 | 1:200 |
| CD8 | EPR20305 | Abcam | 1:100 | Opal480 | 1:200 |
| MHCI (H2Db) | AF6-88.5.5.3 | Thermofisher | 1:100 | Opal690 | 1:200 |
| PanCK | PCK-26 | Abcam | 1:100 | Opal780 | 1:100 |

**Table S4a. List of mouse strains**

| **Gene** | **Genetic** | **Strain** | **References** |
|---|---|---|---|
| *Kras*$^{fl}$ | *LSL-KrasG12D* knock-in | *Kras*$^{tm4Tyj}$ | [2] |
| *Ptf1a*$^{Cre}$ | *Cre* knock-in | *Ptf1a*$^{tm1(cre)Hnak}$ | [3] |
| *Trp53*$^{fl}$ | *LoxP*-sites knock-in | *Trp53*$^{tm1Brn}$ | [4] |
| *Kras*$^{frt}$ | *FSF-KrasG12D* knock-in | *Kras*$^{tm1Dsa}$ | [5] |
| *Ptf1a*$^{Flp}$ | *Flp* knock-in | *Ptf1a*$^{tm(flp)}$ | unpublished |
| *Trp53*$^{frt}$ | *frt*-sites knock-in | *Trp53*$^{tm1.1Dgk}$ | [6] |
| *Rosa26 locus (Cre*$^{ERT2}$*)* | *FSF-Cre*$^{ERT2}$ *knock-in* | *Gt(ROSA)26Sor*$^{tm3(CAG-Cre/ERT2)Das}$ | [5, 7] |
| *Rosa26 locus (GP*$^{f}$*l)* | *LSL-GP knock-in* | *Gt(ROSA)26Sor*$^{tmloxP-STOP-loxP-GP-IRES-YFP}$ | [8] |
| *unknown locus* | *TCR transgene* | *Tg(TcrLCMV)*$^{327Sdz}$ | [9] |

**Table S4b. Interbred mouse strains and description.**

| Strains intercrossed | Strain (abbreviated) | Action | Genotype |
|---|---|---|---|
| *Ptf1a*$^{tm1(cre)Hnak}$<br>*Kras*$^{Tm4Tyj}$<br>*Trp53*$^{tm1Brn}$ | *CKP* | Induction of spontaneous PDAC through CRE-induced pancreas-specific activation of mutant KRAS$^{G12D}$ and homozygous loss of Tp53 | *Ptf1a*$^{wt/Cre}$;*Kras*$^{wt/LSL-G12D}$;*p53*$^{fl/fl}$ |
| *Kras*$^{tm1Dsa}$<br>*Ptf1a*$^{tm(flp)}$<br>*Trp53*$^{tm1.1Dgk}$<br>*Gt(ROSA)26Sor*$^{tm3(CAG-Cre/ERT2)Das}$<br>*Gt(ROSA)26Sor*$^{tmloxP-STOP-loxP-GP-IRES-YFP}$ | *FKPC2GP* | Induction of spontaneous PDAC through Flp-induced pancreas-specific activation of mutant KRAS$^{G12D}$ and homozygous loss of Tp53. Tamoxifen-inducible pancreas-specific expression of YFP and glycoprotein | *Ptf1a*$^{wt/Flp}$;*Kras*$^{wt/FSF-G12D}$;*p53*$^{frt/frt}$;*ROSA26*$^{FSF-CreERT2/LSL-GP}$ |

**Table S5. FACS antibodies**

| Antigen | Conjugate | Clone | Manufacturer | Isotype | Reactivity |
|---|---|---|---|---|---|
| PGRN | Unconjugated | A23 | Ref [1] | Mouse IgG1k | Human, Mouse |
| HLA-A/B/C | FITC | W6/32 | Biolegend | Mouse IgG2a | Human |
| HLA-DR | FITC | L243 | Biolegend | Mouse IgG2a | Human |
| LCMV-gp33 | FITC | -- | In house | | Mouse |
| H2Db | FITC | KH95 | Biolegend | Mouse IgG2b | Mouse |
| CD3 | APC | 145-2C11 | BD | Armenian Hamster IgG1k | Mouse |
| CD45.1 | FITC | A20 | Thermofisher | Mouse IgG2a | Mouse |
| CD8 | eFluor450 | 53-6.7 | ebiosciences | Rat Ig2a | Mouse |
| GzmB | Alexa Fluor 647 | GB11 | Biolegend | Mouse IgG1k | Mouse |
| TNFa | PE | MP6-XT22 | Thermofisher | Rat IgG1k | Mouse |
| IFNg | PE | XMG1.2 | Thermofisher | Rat IgG1k | Mouse |
| EpCAM | APC | G8.8 | Thermofisher | Rat IgG2ak | Mouse |
| $\alpha$-sma | Unconjugated | 1A4 | Thermofisher | Mouse IgG2a | Mouse |

**Table S6. Immunofluorescence staining and antibodies**

| Primary antibodies | Clone | Manufacturer | Dil. | Secondary antibodies | Manufacturer | Dil. |
|---|---|---|---|---|---|---|
| LC3B | Poly | Abcam | 1:500 | Dylight594-Goat anti-rabbit | Thermofisher | 1:500 |
| HLA-A/B/C | W6/32 | Biolegend | 1:150 | Alexa Fluor488-Goat anti-mouse | Thermofisher | 1:500 |
| H2Db | AF6-88.5.5.3 | ThermoFisher | 1:150 | Alexa Fluor488-Goat anti-mouse | Thermofisher | 1:500 |

**Table S8. Association of PGRN level with clinicopathological parameters and immune markers in CONKO-001 cohort**

| | PGRN_Lo | PGRN_Hi | *P* value |
|---|---|---|---|
| **Tumor size_2 classes** | | | |
| T1-2 | 6 | 1 | *0.107* |
| T3-4 | 30 | 34 | |
| **Ki67** | | | |
| <50 | 17 | 9 | *0.067* |
| 50+ | 16 | 22 | |
| **SMA_intensity** | | | |
| Low | 2 | 7 | *0.247* |
| High | 16 | 21 | |
| **P53 intensity** | | | |
| 0-1 | 13 | 9 | *0.292* |
| 2-3 | 19 | 23 | |
| **CD8+** | | | |
| <42 | 9 | 16 | ***0.037*** |
| 42+ | 25 | 15 | |
| **Stromal CD8+** | | | |
| ≤25 | 8 | 12 | *0.185* |
| 25+ | 26 | 19 | |
| **CD103+** | | | |
| ≤3 | 27 | 23 | *0.251* |
| 3+ | 6 | 10 | |

#### REFERENCES

1. Ho JC, Ip YC, Cheung ST, Lee YT, Chan KF, Wong SY, Fan ST: **Granulin-epithelin precursor as a therapeutic target for hepatocellular carcinoma**. *Hepatology* 2008, **47**(5):1524-1532.
2. Jackson EL, Willis N, Mercer K, Bronson RT, Crowley D, Montoya R, Jacks T, Tuveson DA: **Analysis of lung tumor initiation and progression using conditional expression of oncogenic K-ras**. *Genes & development* 2001, **15**(24):3243-3248.
3. Nakhai H, Sel S, Favor J, Mendoza-Torres L, Paulsen F, Duncker GI, Schmid RM: **Ptf1a is essential for the differentiation of GABAergic and glycinergic amacrine cells and horizontal cells in the mouse retina**. *Development* 2007, **134**(6):1151-1160.
4. Marino S, Vooijs M, van Der Gulden H, Jonkers J, Berns A: **Induction of medulloblastomas in p53-null mutant mice by somatic inactivation of Rb in the external granular layer cells of the cerebellum**. *Genes & development* 2000, **14**(8):994-1004.
5. Schonhuber N, Seidler B, Schuck K, Veltkamp C, Schachtler C, Zukowska M, Eser S, Feyerabend TB, Paul MC, Eser P *et al*: **A next-generation dual-recombinase system for time- and host-specific targeting of pancreatic cancer**. *Nature medicine* 2014, **20**(11):1340-1347.
6. Lee CL, Moding EJ, Cuneo KC, Li Y, Sullivan JM, Mao L, Washington I, Jeffords LB, Rodrigues RC, Ma Y *et al*: **p53 functions in endothelial cells to prevent radiation-induced myocardial injury in mice**. *Sci Signal* 2012, **5**(234):ra52.
7. Wen HJ, Gao S, Wang Y, Ray M, Magnuson MA, Wright CVE, Di Magliano MP, Frankel TL, Crawford HC: **Myeloid Cell-Derived HB-EGF Drives Tissue Recovery After Pancreatitis**. *Cell Mol Gastroenterol Hepatol* 2019, **8**(2):173-192.
8. Page N, Klimek B, De Roo M, Steinbach K, Soldati H, Lemeille S, Wagner I, Kreutzfeldt M, Di Liberto G, Vincenti I *et al*: **Expression of the DNA-Binding Factor TOX Promotes the Encephalitogenic Potential of Microbe-Induced Autoreactive CD8(+) T Cells**. *Immunity* 2018, **48**(5):937-950 e938.
9. Pircher H, Burki K, Lang R, Hengartner H, Zinkernagel RM: **Tolerance induction in double specific T-cell receptor transgenic mice varies with antigen**. *Nature* 1989, **342**(6249):559-561.
