## Supplementary Experimental Procedures for "Progranulin promotes immune evasion of pancreatic adenocarcinoma through regulation of MHCI expression"

### Cell culture and treatments

Human PDAC cell lines, PaTu8988T and MiaPaCa2 were purchased from the American Type Culture Collection. Stable cell lines for GRN suppression were established by transfecting *GRN* shRNA into Patu8988T and MiaPaCa2. Scramble shRNA was included as negative control (nc) for transfection.

Sequences of shRNA and nc are as follows,
sh_GRN1:
AGGCCCTGATAGTCAGTTCGAATtgacaggaagATTCGAACTGACTATCAGGGC
sh_GRN2:
AGGAAGGACACTTCTGCCATGATtgacaggaagATCATGGCAGAAGTGTCCTTC
sh_GRN3:
AGGTGACCTGATCCAGAGTAAGTtgacaggaagACTTACTCTGGATCAGGTCAC
nc:
AGGGAATCTCATTCGATGCATACtgacaggaagGTATGCATCGAATGAGATTCC
All transfectants were maintained in 10% FBS-supplemented DMEM with 2 mg/mL of G418 (Life Technologies, Thermo Fisher Scientific, MA).

### Enzyme-linked immunosorbent assay

Soluble PGRN levels in human plasma samples and culture supernatants were detected by a human PGRN ELISA kit (Adipogen Inc.). Plasma samples from normal individuals and PDAC patients were diluted at 1:100 using the diluent provided by the kit, while culture supernatants from *in vitro* experiments were undiluted. While for mouse plasma collected from in vivo antibody treatment experiment, plasma PGRN levels were measured by mouse PGRN ELISA kit (Adipogen). Plasma samples were diluted at 1:5 using the diluent provided by the kit.

### Real-Time quantitative Reverse-Transcription Polymerase Chain Reaction

Real-time quantitative PCR (qPCR) was performed by Roche LightCycler® 480 using LightCycler® 480 SYBR Green I Master Kit (Roche GmbH, Germany). Primers for PGRN (*GRN*) were designed using the NCBI Primer Blast and purchased from Eurofins MWG Operon GmbH, Ebersberg, Germany. The PCR products were designed with a size of ~100 bp. Real-time qPCR experiments were run under 58° C annealing condition and amplification was run for 45 cycles. A melting curve was implemented in each experiment to prove single product amplification. Data was analyzed using ΔCt calculations where GAPDH or GUSB served as housekeeper control for normalization. The amplification efficiency was experimentally determined or assumed as 2 (doubling each cycle). Relative mRNA expression levels compared to housekeeper gene expression (efficiency-ΔCt) were used for visualization.

**Immunofluorescence staining**

Cells were grown on 8-well cell culture chamber slides (Lab-Tek, Waltham, USA). 3 days after treatment with a single $IC_{50}$ dose of trametinib or DMSO control (Sigma-Aldrich, St. Louis, USA), slides were washed with PBS 3 times and then fixed and permeabilized with ice-cold Methanol for 5min. After washing with PBS 3 times, slides were blocked with 10% normal goat serum in PBS for 1h. Cells were incubated with a 1:100 dilution of the corresponding primary antibodies **(Table S6)** in 1% BSA at RT for 1hr. Slides were washed and incubated with the secondary antibody (1:400; Thermo Scientific, Waltham, USA) for 1 h at room temperature in the dark. Afterwards, slides were washed 3 times with PBS, counterstained with DAPI (Vector Laboratories, Burlingame, USA) and scanned with AxioScanner.

**Lysosomal staining**

Lysosomal staining was performed using Cytopainter LysoGreen indicator reagent (Abcam) according to manufacturer's instructions. Briefly, cells were grown to about 70% confluence and then incubated with Cytopainter LysoGreen indicator reagent (Abcam) for 1 h at 37 °C with 5% $CO_2$. Cells were then washed twice in pre-warmed Hank's balanced salt solution (HBSS, Life Technologies), trypsinized with 0.05% trypsin EDTA (Life Technologies) and resuspended in 1x phosphate buffered saline (Life Technologies) for flow cytometry. Lysosomal levels were measured by flow cytometer (BD).
